# MOTL: enhancing multi-omics matrix factorization with transfer learning

**DOI:** 10.1101/2024.03.22.586210

**Authors:** David P. Hirst, Morgane Térézol, Laura Cantini, Paul Villoutreix, Matthieu Vignes, Anaïs Baudot

**Affiliations:** Aix Marseille Univ, INSERM, MMG, Centuri, Marseille, France; Institut Pasteur, Université Paris Cité, CNRS UMR 3738, Machine Learning for Integrative Genomics Group, F-75015 Paris, France; School of Mathematical and Computational Sciences, College of Science, Massey University, Palmerston North, New Zealand; CNRS, Marseille, France; Barcelona Supercomputing Center, Barcelona, Spain

**Keywords:** Matrix Factorization, Dimensionality Reduction, Multi-Omics, Data Integration, Transfer Learning, MOFA

## Abstract

Joint matrix factorization is popular for extracting lower dimensional representations of multi-omics data but loses effectiveness with limited samples. Addressing this limitation, we introduce MOTL (Multi-Omics Transfer Learning), a framework that enhances MOFA (Multi-Omics Factor Analysis) by inferring latent factors for small multi-omics target datasets with respect to those inferred from a large heterogeneous learning dataset. We evaluated MOTL by designing simulated and real data protocols and demonstrated that MOTL improves the factorization of limited-sample multi-omics datasets when compared to factorization without transfer learning. When applied to actual glioblastoma samples, MOTL enhanced delineation of cancer status and subtype.

## 1 Background

Omics data have transformed the study of biology and medicine by enabling high-throughput measurements of the activity and abundance of biological molecules and processes (Conesa and Beck, 2019; Dermitzakis, 2008; Manzoni et al., 2018) In recent years, the fields of biology and medicine have been revolutionized by the increased availability of multi-omics datasets (Conesa and Beck, 2019; Subramanian et al., 2020; Huang et al., 2017). A multiomics dataset is comprised of multiple data matrices, each containing a different type of omics data (e.g., mRNA transcript counts, genomic mutations, DNA methylation prevalence). The integrative analysis of multi-omics data can provide a better understanding of a biological system than that obtained from the analysis of a single omics data matrix, as the complementary information contained in different omics enables a more comprehensive overview of the underlying biological system (Rappoport and Shamir, 2018; Pierre-Jean et al., 2020; Huang et al., 2021; Cantini et al., 2021; Ballard et al., 2024; Baião et al., 2025). Additionally, using multiple omics can reveal insights into relationships between the different biological layers they represent (Huang et al., 2017; Chauvel et al., 2020; Subramanian et al., 2020). Combining omics is also expected to reduce the impact of noise (Rappoport and Shamir, 2018; Chauvel et al., 2020; Cantini et al., 2021). However, multi-omics data poses further analysis challenges beyond those encountered in single omics data analysis. These challenges include increased dimensionality, the presence of multiple data types, diverse sources of technological noise, and diverse ranges of variability. In this context, there has been an increased need for methods able to carry out integrative analysis of multiple omics.

The development of multi-omics analysis tools is an active area of research and a large variety of strategies have been proposed. These strategies encompass broad and overlapping categories, including Bayesian methods, network-based approaches, or dimensionality reduction techniques (Subramanian et al., 2020; Cantini et al., 2021), alongside more recent deep learning strategies (Ballard et al., 2024; Baião et al., 2025; Wen et al., 2023). A category of multi-omics analysis tools that is widely used is dimensionality reduction with matrix factorization. Matrix factorization infers a lower dimensional representation of the observed data, in which a sufficiently informative proportion of the original signal is retained (Stein-O’Brien et al., 2018). It has proven to be computationally efficient, but also interpretable, and effective for the analysis of large datasets (Rappoport and Shamir, 2018; Tini et al., 2019; Chauvel et al., 2020; Pierre-Jean et al., 2020; Cantini et al., 2021).

Most classical matrix factorization approaches were designed for the analysis of a single data matrix. Applying matrix factorization to a single omics matrix produces a score matrix and a weight matrix, both of which contain values for latent factors that are potentially associated with different sources of underlying biological signal. The values in the weight matrix ideally represent signal across the assayed biological features, and the values in the score matrix represent the signal across the samples. For a multiomics dataset, one of the strategies is to jointly factorize multiple omics data matrices. Various methods are now available for this purpose (Cantini et al., 2021). Multi-omics joint matrix factorization methods typically produce a weight matrix for each omics, and either a shared score matrix or a combination of shared and omics specific score matrices. Many multi-omics matrix factorization methods are extensions of classical methods. For example, intNMF (Chalise and Fridley, 2017) extends non-negative matrix factorization to the multi-omics setting, and allows the user to determine the relative contribution of each omics to the extraction of joint signal. JIVE (Lock et al., 2013) extends principal component analysis to model both joint and omics specific signal. moCluster (Meng et al., 2016) extends canonical correlation analysis to produce a shared score matrix based on score matrices produced for each of two or more omics. MOFA (Argelaguet et al., 2018), which is an extension of Factor Analysis, uses a Bayesian framework to account for the presence of multiple data types and to distinguish between joint and omics specific signal. Overall, the factors inferred by multi-omics matrix factorization can be used for clustering samples to reveal disease sub-types, for identifying molecular profiles and biomarkers associated with diseases, as well as for prediction of outcomes such as drug response and survival (Rappoport and Shamir, 2018; Taroni et al., 2019; Pierre-Jean et al., 2020; Cantini et al., 2021; Banerjee et al., 2023). A challenge for matrix factorization is that it requires a large amount of observed data to produce a meaningful representation. However, there are cases where omics are measured from only a small number of samples, due to the rareness or cost of obtaining the data, and so there is a need for methods which help mitigate this challenge (Weiss et al., 2016; Stein-O’Brien et al., 2018; Banerjee et al., 2023).

For a dataset generated from a small number of samples, transfer learning is a potential solution to the limited effectiveness of matrix factorization. Transfer learning is a machine learning approach in which information extracted from a large learning domain is used to improve the performance of a task applied to a smaller target domain (Weiss et al., 2016; Stein-O’Brien et al., 2019; Taroni et al., 2019; Banerjee et al., 2023). It is assumed that the two domains share an overlapping latent space, allowing knowledge from the learning domain to be transferred to the application of the task to the target domain. Transfer learning has been successfully used in various machine learning applications, including image classification, text sentiment classification and recommendation systems (Weiss et al., 2016; Veeramachaneni et al., 2019; Dong et al., 2021; Banerjee et al., 2023). In a transfer learning approach to omics matrix factorization, information inferred from the prior factorization of a learning dataset, comprised of a large number of samples from a heterogeneous set of biological conditions, is incorporated into the factorization of a small target dataset (Peng et al., 2021; Banerjee et al., 2023). It is assumed that if the latent factors inferred from the learning dataset represent common underlying biological processes, they should help improve the factorization of the target dataset (Stein-O’Brien et al., 2019).

The usefulness of transfer learning approaches to matrix factorization, for omics data analysis, has been demonstrated in contexts in which both the target and learning datasets were comprised of single matrices of omics data. In these cases, transfer learning was used to infer a score matrix for the target dataset by projecting it onto a weight matrix inferred from a learning dataset. In one study, Stein-O’Brien et al. factorized a mouse single cell RNA-seq learning dataset with the Bayesian non-negative matrix factorization algorithm CoGAPS (Fertig et al., 2010). Then, they used the transfer learning tool projectR (Sharma et al., 2020) to infer a score matrix for a human time course bulk RNA-seq dataset. The resulting factors were associated with known spatiotemporal differences across the samples. In another example, Davis-Marcisak et al. factorized a mouse single cell RNA-seq learning dataset with CoGAPS, and then used projectR to infer a score matrix for bulk RNA-seq data from human cancer samples. They observed an association between a particular projectR factor and outcomes in metastatic melanoma. Taroni et al. developed MultiPLIER, a transfer learning framework, which they demonstrated by firstly applying the non-negative matrix factorization algorithm PLIER (Mao et al., 2019) to a subset of Recount2 to infer a weight matrix. Recount2 is a compendium of RNA-seq data obtained from 70,000+ human samples taken across more than 2,000 studies (Collado-Torres et al., 2017). Taroni et al. then used a blood cell compendium of microarray gene expression data as the target dataset, for which they inferred a score matrix with MultiPLIER, as well as factorizing the target dataset directly with PLIER. For counts of a cell type of interest, MultiPLIER inferred a more highly correlated factor than was inferred by direct factorization of the target dataset. They also used microarray gene expression data for 79 samples from a rare disease group called antineutrophil cytoplasmic autoantibody associated vasculitis (AAV) as a target dataset. There are no AAV samples in the Recount2 compendium, yet the MultiPLIER factors were positively correlated with their best match from factors inferred by direct factorization of the target dataset.

It has thus been demonstrated that the application of matrix factorization to a large, heterogeneous learning dataset can yield factors containing transferable information, that are biologically relevant to target datasets from different organisms, diseases, cell types and omics platforms. However, existing transfer learning approaches to matrix factorization have been designed for, and demonstrated on, datasets comprised of single omics data only. To the best of our knowledge, transfer learning approaches to joint multi-omics matrix factorization are currently lacking.

We here introduce MOTL (Multi-Omics Transfer Learning), a novel Bayesian transfer learning algorithm for multi-omics matrix factorization. MOTL is based on MOFA, a popular tool for integrative multi-omics analysis (Argelaguet et al., 2018). We first present the statistical framework and implementation of MOTL. Next, we propose two protocols, that we designed based on simulated and real multi-omics datasets, for evaluating the performance of transfer learning approaches. We used these protocols to evaluate MOTL, and observed that, for a target multi-omics dataset comprised of a small number of samples, our transfer learning approach to matrix factorization is more effective than matrix factorization without transfer learning. Lastly, we showcase a practical use case of MOTL on a limited glioblastoma sample set, revealing an enhanced delineation of cancer status and subtype thanks to transfer learning.

## 2 Results

### 2.1 MOTL: A new transfer learning framework for multi-omics matrix factorization

We propose MOTL, a transfer learning approach to multi-omics matrix factorization. MOTL is based on MOFA (Argelaguet et al., 2018), which uses variational Bayesian inference (Blei et al., 2017). Consider a multi-omics target dataset, ***T***, consisting of omics matrices, ***T*** ^(*m*)^, *m* = 1, …, *M*. Each 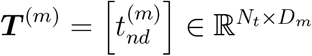 contains data for *N*_*t*_ samples (rows) and *D*_*m*_ features (columns), where 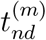 is the value for the *n*th sample and the *d*th feature from the *m*th matrix (See Methods 5.1 for a summary of the mathematical notation used in this document). The features depend on which molecules were assayed to generate a given omics matrix; for example the features for mRNA counts are genes, while those for DNA methylation are CpG sites.

We wish to jointly factorize ***T*** ^(*m*)^ into a matrix of sample scores, 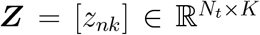, and an omics specific matrix of feature weights, 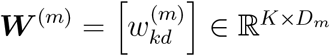. The resulting lower dimensional representation is based on *K* factors, which ideally represent underlying biological signals associated with some biological condition(s) of interest. *z*_*nk*_ is the score for the *n*th sample and the *k*th factor, while 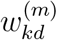 is the weight for the *k*th factor and the *d*th feature from the *m*th matrix. The *k*th column vector of ***Z***, denoted by ***z***_:*k*_, contains scores for factor *k*, while the *n*th row vector, ***z***_*n*:_, contains scores for sample *n*. The *k*th row vector of ***W*** ^(*m*)^, denoted by 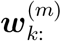, contains weights for factor *k*, for the *m*th matrix, while the *d*th column vector, 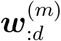, contains weights for feature *d* from the *m*th matrix.

We are concerned with the situation in which *N*_*t*_ is small, exacerbating the curse of dimensionality, and therefore we expect to improve the factorization of ***T*** by employing a transfer learning approach (see Figure 1). We do this transfer learning by incorporating values that have already been inferred from the prior factorization of a learning dataset, ***L***, and we assume that the Bayesian matrix factorization algorithm MOFA (Argelaguet et al., 2018) was used for factorizing ***L***.

**Figure 1:**
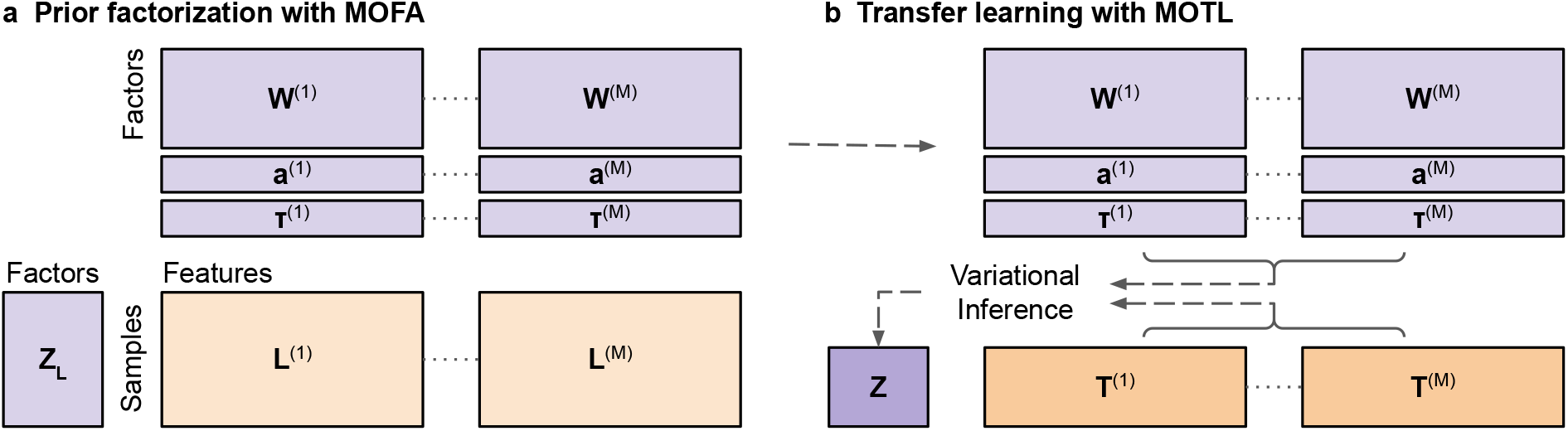
Overview of MOTL, our transfer learning approach to joint multi-omics matrix factorization based on variational Bayesian inference. **a** A multi-omics learning dataset, ***L***, consisting of *M* omics matrices, ***L***^(*m*)^, *m* = 1, …, *M*, is factorized with MOFA to infer a matrix of feature weights, ***W*** ^(*m*)^, vector of feature-wise intercepts, ***a***^(*m*)^, and a vector of feature-wise precision parameter values, ***τ*** ^(*m*)^, for each ***L***^(*m*)^. **b** The feature weight, intercept, and precision parameter values, inferred from the factorization of ***L***, are incorporated into the factorization of a multi-omics target dataset, ***T***, for which MOTL infers a matrix of sample scores, ***Z***, with variational inference.

The learning dataset consists of omics matrices ***L***^(*m*)^, *m* = 1, …, *M*. Each 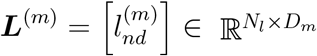 contains data for the same *D*_*m*_ features as ***T*** ^(*m*)^, but for a different set of *N*_*l*_ *> N*_*t*_ samples. We hypothesise that if ***L*** is comprised of samples from a heterogeneous set of biological conditions, then the factorization of ***L*** will yield information that is relevant for the factorization of ***T***.

MOTL is based on the variational Bayesian inference methodology used by MOFA (Methods 5.2). We have modified the MOFA algorithm to enable us to supplement the factorization of ***T*** by incorporating values already inferred from the prior factorization of ***L***. For MOTL, we assume that each observed 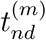 is a random variable, with a likelihood that is conditional on vectors ***z***_*n*:_ and 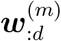. We model continuous, counts and binary data with the same likelihoods and link functions that MOFA uses. For observed continuous data we thus assume a Gaussian likelihood, into which we include a feature-wise precision parameter, 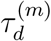, for each feature *d* from matrix *m*. For observed binary data we assume a Bernoulli likelihood, and for observed counts data we assume a Poisson likelihood. In contrast to MOFA, MOTL doesn’t center the input data during factorization fitting, as we want to incorporate an intercept that is compatible with the factorization of ***L***. We therefore replace 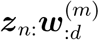 with 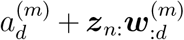 in the likelihood, where 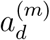 is the feature-wise intercept for feature *d*, from matrix *m*. We infer 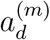 values based on the MOFA factorization of ***L*** (Methods 5.7). MOTL accepts missing 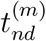 values; therefore it is not necessary to remove features with missing values, or perform imputation, before using MOTL.

In order to carry out a transfer learning approach to matrix factorization, MOTL uses the matrix of feature weights, ***W*** ^(*m*)^, vector of feature-wise intercepts, 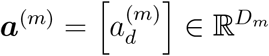, and vector of feature-wise precision parameter values, 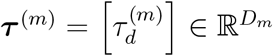, inferred for each ***L***^(*m*)^ with a prior MOFA factorization of ***L***. Instead of modelling these as random variables, we treat them as constants. We aim to obtain point estimates of *z*_*nk*_ values, for which we assume the same joint prior distribution as MOFA does,

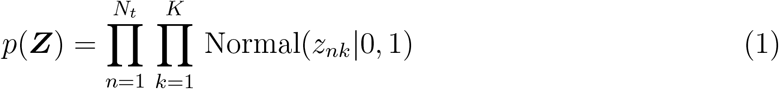

MOTL obtains point estimates of *z*_*nk*_ values by approximating the joint posterior distribution *p*(***Z***|***T***) with a variational distribution:

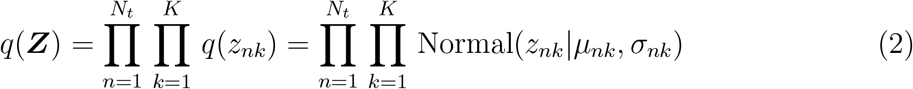

MOTL infers *q*(***Z***) iteratively. At each iteration, the value of each parameter is updated while all other parameter values are held fixed. MOTL optimizes the joint variational distribution by iterating until convergence. For each *z*_*nk*_, the expected value, 𝔼_*q*_ [*z*_*nk*_] = *µ*_*nk*_, is used as the point estimate throughout and after model fitting. MOTL uses the same update equations for the parameters of *q*(*z*_*nk*_) as MOFA, but with the inclusion of intercepts:

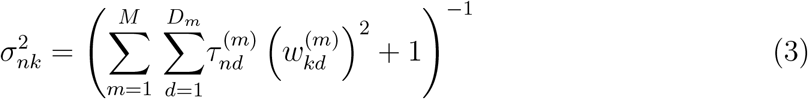

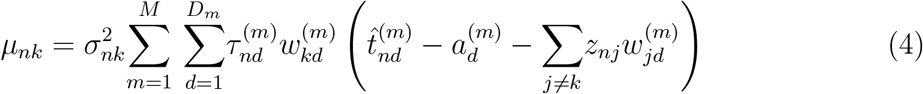

where 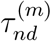 is the precision for the *n*th sample and *d*th feature from the *m*th matrix, and 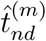 denotes a (possibly) transformed observed data point (Methods 5.2). For observed data with a Gaussian assumed likelihood, a feature-wise precision, 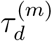, is used instead of 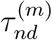, and there is no transformation, meaning 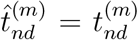. For observed data with a non-Gaussian assumed likelihood, MOTL transforms the data to yield Gaussian pseudo-data values, which it does not center. The transformation to Gaussian pseudo-data allows updates of *q*(***Z***) to be based on the assumption of Gaussian observed data. When MOFA transforms observed data with a Bernoulli assumed likelihood, it derives and uses a precision parameter, 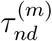, for each sample and feature. For observed data with a Poisson assumed likelihood, it derives and uses a feature-wise precision, 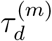. Thus for Bernoulli observed data, MOTL initializes 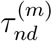 values with 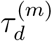 values, which are averages of the 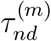 values returned by the factorization of ***L***, and these are subsequently updated at each iteration of the algorithm. For Poisson observed data, MOTL uses the 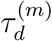 values obtained from the prior factorization of ***L***, and holds them fixed.

To monitor convergence we calculate the evidence lower bound (ELBO), which can be used to evaluate how well a variational distribution approximates a posterior distribution of interest. We calculate the ELBO with respect to ***Z***:

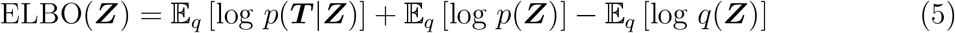

For ***T*** ^(*m*)^ with a non-Gaussian assumed likelihood, we use the same lower bound for 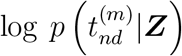 as MOFA does. Maximizing this lower bound, coupled with the use of 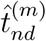 values, allows updates of *q*(***Z***) based on the assumption of Gaussian observed data (Jaakkola and Jordan, 2000; Seeger and Bouchard, 2012). We calculate the ELBO at regular intervals, and the number of iterations between each calculation is a user defined parameter. We check for convergence based on the absolute change in the ELBO (from the previous check) as a percentage of the initial ELBO. The algorithm is deemed to have converged when a specified number of changes in the ELBO are consecutively below a threshold. Both the threshold, and the required number of consecutive changes falling below this threshold, are user defined parameters.

We allow factors to be dropped during training, based on the fraction of variance explained:

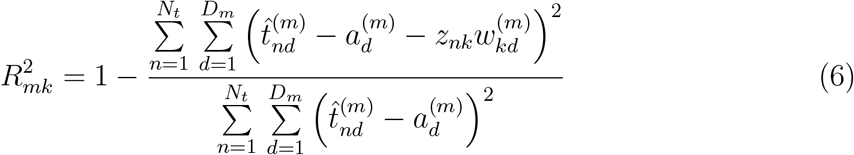

We drop the factor with the lowest 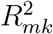 that does not have any 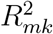 above the threshold. We assess factors in this way after each round of updates. After convergence the algorithm returns ***Z*** and ***W*** ^(*m*)^ matrices for the factors that have not been dropped.

MOTL is available as an open source R implementation (Section 6.3)

### 2.2 Evaluation protocol using simulated multi-omics data

We first designed and implemented a transfer learning evaluation protocol based on simulated multi-omics datasets, which we generated from groundtruth factors (Methods 5.3). In each simulation instance, we generated a multi-omics dataset, ***Y***, which we subsequently split into a target dataset, ***T***, and a learning dataset, ***L. Y*** consisted of matrices of counts, continuous, and binary data. We generated each matrix, ***Y*** ^(*m*)^, from a statistical distribution conditional on random matrices ***Z*** and ***W*** ^(*m*)^, which each contained values for *K* groundtruth factors. The *k*th column vector of ***Z*** contained sample scores for the *k*th groundtruth factor. The *k*th row vector of ***W*** ^(*m*)^ contained feature weights for that same factor. We varied the number of groundtruth factors across configurations, using *K* ∈ {20, 30}. We generated ***Z*** based on the group membership of samples. In each instance, we created two groups of five samples for the target dataset. The learning dataset samples belonged to either 20 or 40 differently sized groups of randomly selected sizes. For each groundtruth factor and group, the sample scores were generated using a mean parameter value that was common to all samples in the group. We induced heterogeneity by allowing the means to vary across groups and factors, randomly selecting each group mean, for a given groundtruth factor, from a pool of three possible values. We split each ***Y*** ^(*m*)^ into ***T*** ^(*m*)^ and ***L***^(*m*)^, based on the sample groups used to generate ***Z***. In each instance ***T*** contained data for 10 samples, while the expected number of samples for ***L*** was ∈ {400, 1000}.

For each simulation instance, we factorized ***L*** with MOFA (Methods 5.6). We then factorized ***T*** with our transfer learning method MOTL (Methods 5.7), incorporating output from the factorization of ***L***. To benchmark the performance of MOTL, we also performed direct MOFA factorizations (i.e., factorization without transfer learning) of ***T*** datasets. We evaluated both the MOTL and direct MOFA factorization of each ***T***, and compared the overall performance of each approach. We evaluated factorizations of each ***T*** by calculating an F1 score (Methods 5.8), to measure how well the factorization allowed us to uncover differentially active groundtruth factors underlying ***T***. The *k*th groundtruth factor was differentially active for ***T*** if the mean parameter values used to simulate the sample scores, for that factor, differed between the two groups of target dataset samples. Factorization with MOTL led to higher F1 scores than direct MOFA factorization, indicating that the MOTL factorizations were more effective in uncovering differentially active latent signal from ***T*** datasets (Figure 2**a**). This was observed across all simulation configurations, and the overall uplift in mean F1 score for MOTL, when compared to direct MOFA factorization, was 0.21 (p-value *<* 0.01, Methods 5.8). We thus observed that transfer learning with MOTL was more effective in uncovering differentially active latent signal, when compared to direct MOFA factorization (without transfer learning) of ***T***. Of note, MOTL was also more effective than direct factorization with the alternative multi-omics matrix factorization approaches intNMF and moCluster (Supplementary Figure S1).

**Figure 2:**
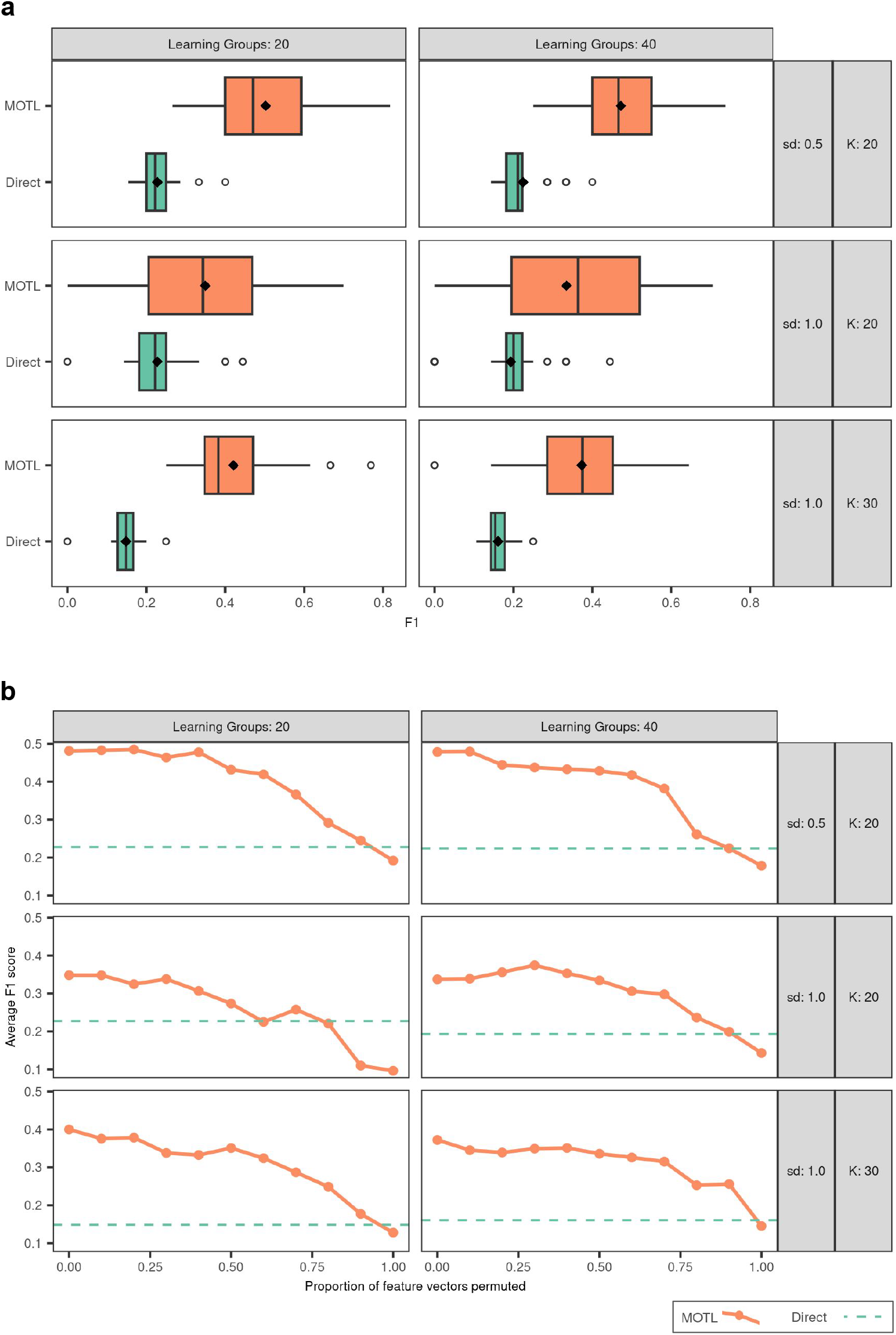
Evaluation of factorizations of small simulated multi-omics target datasets. **a** The boxplots represent the F1 scores obtained for factorizations with and without MOTL transfer learning, for different simulation configuration settings. Simulation configurations varied in the number of groups of samples used for the learning dataset (*Learning Groups*), the number of groundtruth factors (*K*), or the standard deviation used to simulate *z*_*nk*_ values (*sd*). F1 scores take a value between 0 and 1, and higher values indicate better factorizations. Each boxplot is based on 30 F1 scores. The hinges of the boxes are the 25^*th*^ and 75^*th*^ percentiles, the middle lines are medians, the diamonds are the mean values, and the whiskers are either extreme values or extend 1.5 times the inter-quartile range from the hinge. **b** The line plots represent the average F1 score obtained from MOTL factorizations, for different simulation configuration settings, after permuting the values in proportions of the feature vectors in the ***W*** ^(*m*)^ matrices obtained from prior factorization of ***L*.** The dashed line is the average F1 score obtained with direct MOFA factorization for the simulation configuration.

We next wanted to evaluate the robustness of MOTL when there is a decline in the overlap between the latent spaces of ***L*** and ***T*** (Methods 5.7). We forced this decline in overlap by permuting feature vectors in the ***W*** ^(*m*)^ matrices inferred from ***L*** datasets, based on a range of permutation proportions between 0 and 1. For each simulation instance, and permutation proportion, *p*, we created new ***W*** ^(*m*)^ matrices by permuting the values in (*p* × 100)% of the feature vectors in the ***W*** ^(*m*)^ matrices inferred from ***L***. We then factorized ***T*** with MOTL, using the permuted ***W*** ^(*m*)^ matrices, and calculated the F1 score. We observed that MOTL outperformed direct MOFA factorization even when there were large declines in the overlap between the latent spaces of ***L*** and ***T***, and that the performance of MOTL tended to drop below that of direct MOFA factorization when the values in 80% or more of the feature vectors were permuted (Figure 2**b**).

### 2.3 Evaluation protocol using TCGA multi-omics data

We next designed, and implemented, a second transfer learning evaluation protocol, based on TCGA multi-omics data (Methods 5.4). We used four types of omics data: log2 transformed mRNA counts, log2 transformed miRNA counts, DNA methylation M-values, and simple nucleotide variation (SNV) binary data, which we obtained for 32 different cancer types. We created target datasets using data from three cancer types; acute myeloid leukemia (LAML), pancreatic adenocarcinoma (PAAD) and skin cutaneous melanoma (SKCM). We created these target datasets by firstly creating four reference datasets. Each reference dataset, ***R***, contained multi-omics data for all samples from either two, or all three of the cancer types. We then randomly split every ***R*** into non-overlapping target datasets which each contained only five samples per cancer type (Figure 3**a**). We merged data from the remaining 29 cancer types into a learning dataset, ***L***, which contained multi-omics data for 7,217 samples.

**Figure 3:**
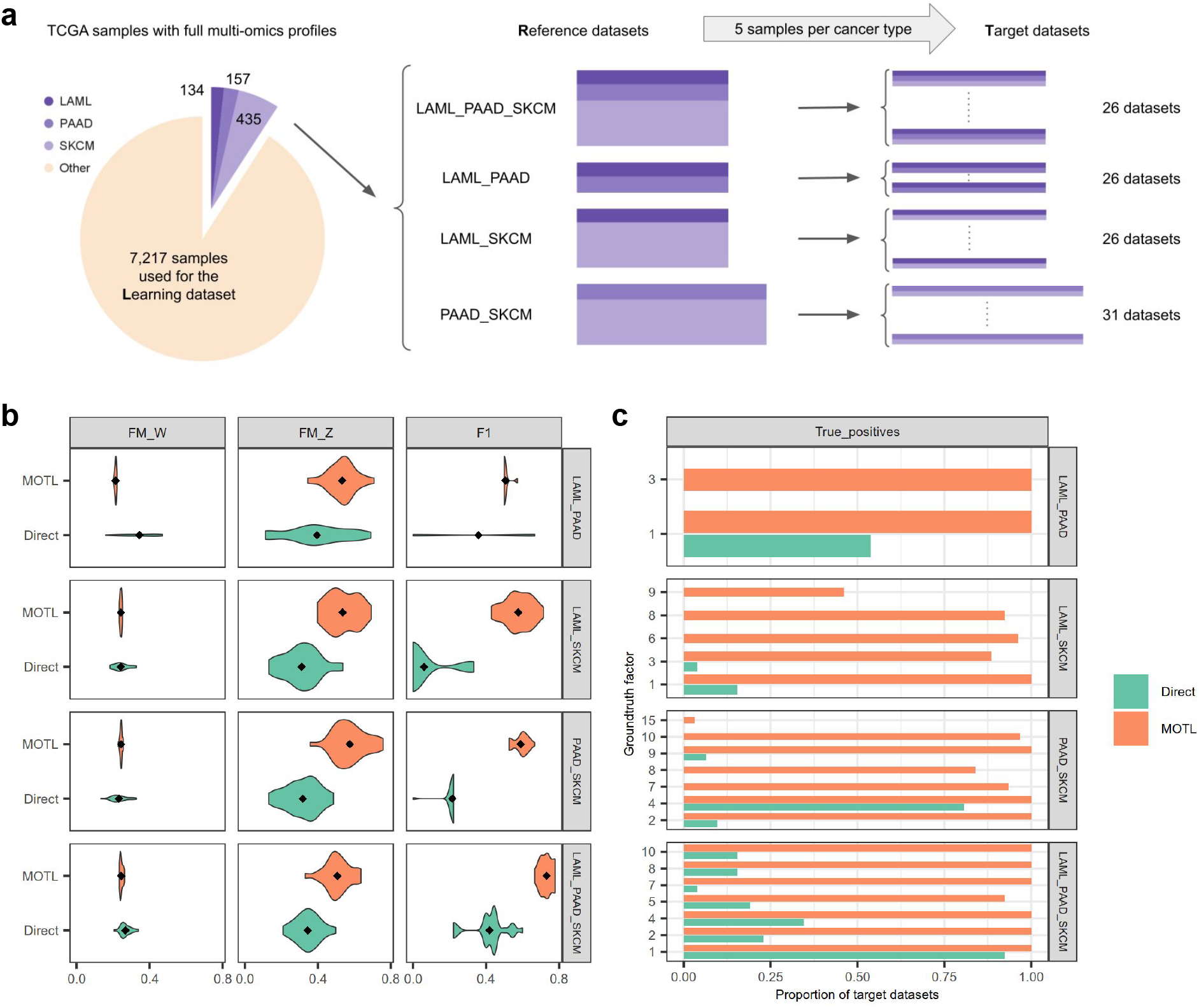
Evaluations using TCGA multi-omics data. **a** We created TCGA target, ***T***, datasets from two or three cancer types: For each combination of cancer types we created a reference, ***R***, dataset containing multi-omics data for all samples from the selected cancer types. We then randomly split ***R*** into non-overlapping ***T*** datasets, containing multi-omics data for five samples per cancer type. We did this for subsets of the set of cancer types {LAML, PAAD, SKCM}. In total we created, and split, four reference datasets, each of which contained multi-omics data for all samples from either two (LAML and PAAD, LAML and SKCM, PAAD and SKCM), or three (LAML, PAAD and SKCM) cancer types. For datasets containing PAAD and SKCM samples only, we included mRNA, miRNA, DNA methylation, and SNV data. We did not include SNV data in datasets containing LAML samples, due to the sparsity of SNV data for LAML. **b** Comparison of factorization approaches applied to TCGA multi-omics datasets. Violin plots of F-measure values for weight matrix factors (*FM W*), F-measure values for score matrix factors (*FM Z*), and F1 scores (*F1*). For each evaluation score, higher values indicate better factorizations. Scores are plotted by factorization method and by the cancer types characterizing the target dataset samples. **c** Frequency with which differentially active groundtruth TCGA factors were true positives. A differentially active groundtruth factor was a true positive if it was predicted as being differentially active based on a factorization of a target dataset. Each bar represents the proportion of target datasets, for which the factorization led to the differentially active groundtruth factor being a true positive. Proportions are plotted by factorization method, and by the cancer types characterizing the reference and target dataset samples.

We factorized ***L*** with MOFA (Methods 5.6), based on which we used MOTL to factorize each ***T*** (Methods 5.7). To benchmark the performance of MOTL, we also performed direct MOFA factorizations (without transfer learning) of ***T*** datasets. In order to evaluate the factorizations of ***T*** datasets, we factorized ***R*** datasets with MOFA and treated the resulting score, ***Z***, and weight, ***W*** ^(*m*)^, matrices as groundtruth factor matrices (Methods 5.8). We were interested in how well the factorizations of ***T*** datasets uncovered the groundtruth, and we used F-measure values and F1 scores to evaluate this.

We calculated F-measure values to assess the correlation between factors inferred from each ***T***, and the groundtruth factors obtained from the factorization of the corresponding ***R*** dataset (Methods 5.8). We calculated F-measure values for weight matrices (FM W), as well as for score matrices (FM Z). The overall mean FM W for MOTL was slightly lower (0.03 reduction, p-value *<* 0.01) than for direct MOFA factorizations of ***T*** datasets (Figure 3**b**, column 1), which is the result of lower average relevance counterbalancing higher average recovery. We concluded from this that groundtruth ***W*** ^(*m*)^ factors were more easily uncovered with those transferred from ***L*** than with direct MOFA factorization. However, despite factor trimming during MOTL factorization, some remaining transferred factors were less associated with groundtruth factors than those obtained with direct MOFA factorization. It is of note that the difference in average FM W is attributable to the datasets containing LAML and PAAD samples only. If we exclude these, there is no difference in FM W (p-value 0.59). The overall mean FM Z for MOTL was 0.20 higher (p-value *<* 0.01, Methods 5.8) than for direct MOFA factorizations (Figure 3**b**, column 2). We thus observed that the ***Z*** factors obtained with MOTL, from ***T*** datasets, were more correlated with groundtruth factors, overall, than those obtained with direct MOFA factorization.

We also calculated F1 scores to measure how well the factorizations of ***T*** datasets uncovered differentially active groundtruth factors (Methods 5.8). For each ***T*** dataset, the groundtruth factors were the factors obtained from factorization of the corresponding ***R*** dataset. We considered the *k*th groundtruth factor to be differentially active if the distribution of scores in the *k*th column vector, of groundtruth ***Z***, differed between the cancer types. We can simultaneously evaluate the ***Z*** and ***W*** ^(*m*)^ factors, and assess the overall quality of factorizations, by checking for an uplift in F1 scores (Figure 3**b**, column 3). MOTL (0.34 uplift, p-value *<* 0.01) yielded higher F1 scores than direct MOFA factorization, meaning it was more effective in uncovering latent activity that varied across cancer types. Similarly to the evaluation using simulated data, MOTL was also more effective than direct factorization with the alternative multi-omics matrix factorization approaches intNMF and moCluster (Supplementary Figure S2).

We next examined differentially active groundtruth factors, with an initial focus on the frequency with which these factors were true positives (Figure 3**c**). For each factorization of a ***T*** dataset, a differentially active groundtruth factor was a true positive if it was predicted as being differentially active based on the factorization of ***T***. The unique count of true positives was a component of each F1 score value (Methods 5.8). We further performed a gene set enrichment analysis to identify the pathways and processes associated with differentially active groundtruth factors that were true positives (Methods 5.8, Supplementary Table 1), and that explained at least 1% of the mRNA variance in ***R*** (Supplementary Table 2).

The factorization of the ***R*** dataset containing all LAML and PAAD samples yielded six groundtruth factors, of which two were differentially active; Factor 1 and Factor 3. Both of these factors were true positives for 100% of MOTL factorizations of ***T*** datasets containing subsets of five LAML and PAAD samples (Figure 3**c**). In contrast, only one of these factors was a true positive for direct MOFA factorizations, and for just over half of the same ***T*** datasets (Figure 3**c**). Factor 1 is significantly associated with developmental processes, cell communication and immunity signaling. Factor 3 displays similar enrichments, with an additional specific enrichment related to the regulation of gene expression in beta cells (Supplementary Table 1).

The factorization of the ***R*** dataset containing all LAML and SKCM samples yielded 12 groundtruth factors, of which five were differentially active. Four out these five factors were true positives for MOTL factorizations for more than 80% of the ***T*** datasets; one factor was a true positive for just under half of the ***T*** datasets. Importantly, only two of these five groundtruth factors were true positives for direct MOFA factorizations, and only for a small proportion of the ***T*** datasets. Four out of these five differentially active groundtruth factors, that were true positives, explained at least 1% of the mRNA variance in ***R***: Factor 1, Factor 3, Factor 8, and Factor 9. We performed gene set enrichment analysis on these four factors. Factor 1 is significantly associated with extracellular organisation, developmental processes, cell communication signaling, and Fc Receptor mediated immune processes (Supplementary Table 1). Factor 3 is significantly associated with hematopoeitic cell lineage, Pi3K/AKT and G protein-coupled signaling, and chemokine, interleukin, interferon signaling. Both Factors 1 and 3 are associated with keratinisation and formation of the cornified envelope. Factors 8 and 9 do not present significant pathway enrichments beyond keratinisation and processes already associated with the first two factors.

The factorization of the ***R*** dataset containing all PAAD and SKCM samples yielded 17 groundtruth factors, of which eight were differentially active. Seven of these eight factors were true positives for MOTL factorizations, six with high frequency (i.e., identified for more than 80% of the ***T*** target datasets). Only one differentially active groundtruth factor is frequently a true positive for direct MOFA factorization of the ***T*** target datasets. Factor 2 is related to B cell receptor signaling and Fc Receptor mediated immune processes, drug metabolism by cytochrome p450 and other metabolism-related processes. This Factor 2 is rarely uncovered by direct MOFA factorization. Contrarily, Factor 4 is a true positive for MOTL for 100% of the target datasets and for direct MOFA factorizations for more than 75% of the target datasets. This factor is associated with developmental processes and cytokine-cytokine receptor interactions. Factor 7 is associated with keratinisation and formation of the cornified envelope, Factor 9 with cell adhesion and migration, and Factor 10 with cytokine and chemokine signaling.

The factorization of the ***R*** dataset containing all LAML, PAAD and SKCM samples yielded 13 groundtruth factors, of which seven were differentially active. All seven of these differentially active groundtruth factors were true positives for MOTL factorization with high frequency, compared to one factor for direct MOFA factorization. These groundtruth factors, differentially active between all three cancer types, are associated with the same cellular processes and pathways identified when comparing the cancer types pairwise (Supplementary Table 1).

Overall, the factors that were differentially active when comparing two or the three cancer types reflect the different embryonic origins of the cancerous tissues, and highlight the importance of immunity and microenviroment in cancer pathophysiology and response to treatments. In conclusion, matrix factorization with transfer learning using MOTL better uncovers differentially active groundtruth factors from target datasets containing only a small number of samples.

### 2.4 Application of MOTL to glioblastoma

Glioblastoma is a rare, heterogeneous, and aggressive cancer type. Multi-omics datasets offer an important opportunity to better characterize glioblastoma subtypes, identify biomarkers, and propose novel therapeutic options (Santamarina-Ojeda et al., 2023). However, large collections of glioblastoma tissue samples are difficult to obtain due to the relative scarcity of the disease and the challenges involved in acquiring samples via invasive biopsies.

In Santamarina-Ojeda et al. (2023), the authors conducted a multi-omics profiling (mRNA expression, DNA methylation) for four normal brain samples and nine patient-derived glioblastoma stem cell (pd-GBSC) cultures. The nine cancer samples had been previously classified into three subtypes thanks to transcriptome-based signatures: classical (CL), proneural (PN), and mesenchymal (MS). Given the small number of samples, the authors devised a strategy based on analyzing this dataset in parallel with datasets gathered from the literature. We illustrate here how MOTL could help in analysing such a dataset comprised of a limited number of samples.

We first applied a direct MOFA factorization (Methods 5.6) to a target dataset comprised of the four normal and nine pd-GBSC samples (Methods 5.5), revealing eight factors. Heatmap clustering of the samples, based on these factors, does not demonstrate clear grouping with respect to either cancer status or subtype (Figure 4a). Next, we applied MOTL to the same target dataset (Methods 5.7). In this case, we first created a TCGA learning dataset containing mRNA expression, miRNA expression, DNA methylation, and SNV data for samples from all 32 cancer types (Methods 5.4). It is noteworthy that this learning dataset did not contain data for glioblastoma, as there were no TCGA glioblastoma samples fulfilling our selection criteria (i.e., with complete 4-layer multi-omics profiles). We factorized this learning dataset with MOFA (Methods 5.6), based on which we applied MOTL to the target dataset. MOTL transfer learning factorization revealed 25 factors. In this case, the heatmap clustering of the MOTL factors separates cancer and normal samples and also displays subgroups partially matching subtypes previously defined based on transcriptome signatures (Figure 4**b**).

**Figure 4:**
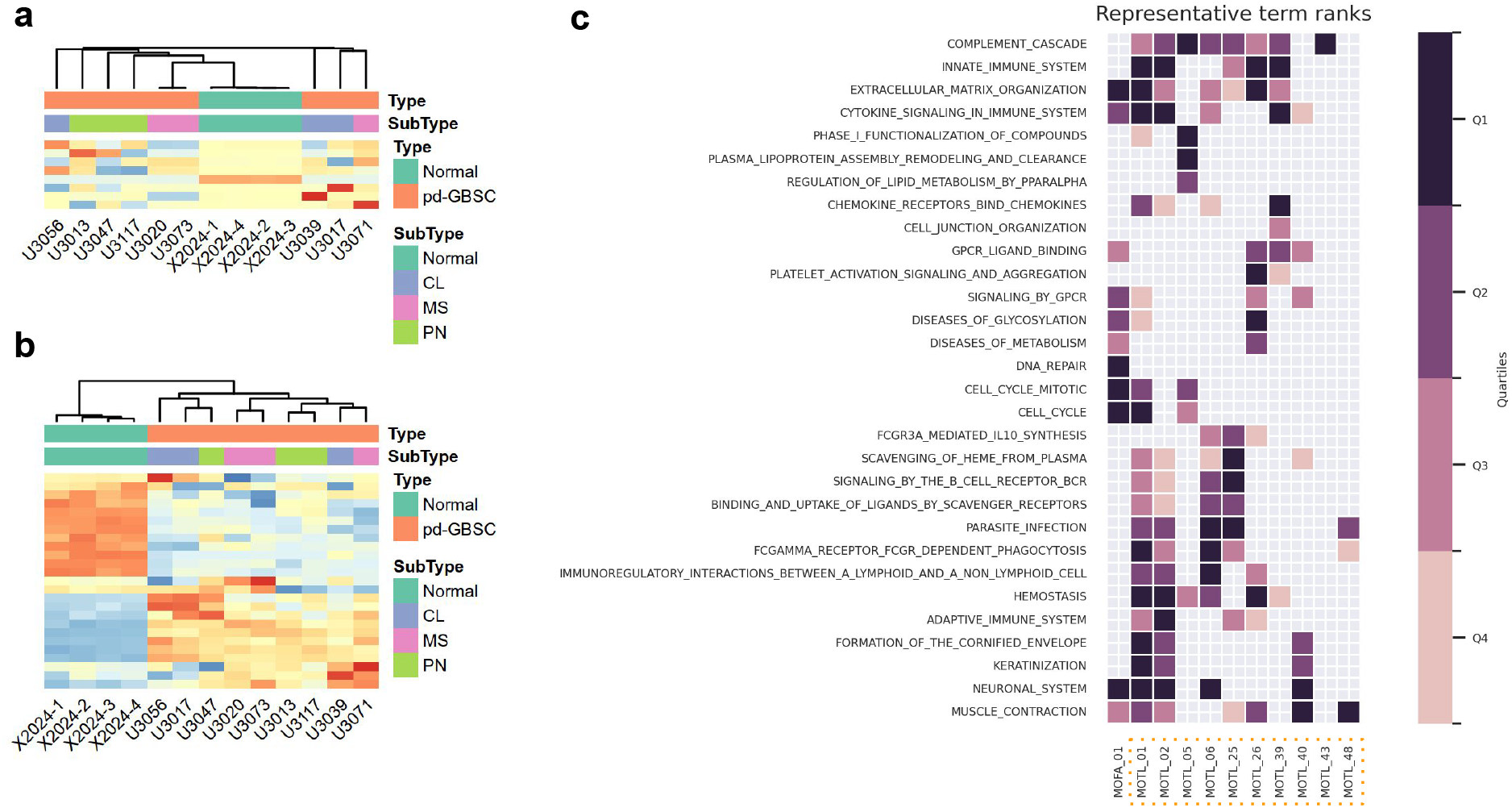
Heatmaps of factorizations and gene set enrichment analysis of glioblastoma data: The heatmaps are based on factorizations and subsequent gene set enrichment analysis of the target dataset comprised of multi-omics data (mRNA expression, DNA methylation) for all normal and patient-derived glioblastoma stem cell (pd-GBSC) samples. The multi-omics data was obtained from Santamarina-Ojeda et al., who also provided subtypes for the cancer samples, previously defined from transcriptomic signatures: CN (classical), PN (proneural), MS (mesenchymal). **a** Heatmap of the score matrix, ***Z***, inferred with Direct MOFA factorization of the target dataset. The rows are the factors, the columns are the samples, and the cells are the row-wise centered and scaled factor values. The rows and columns were ordered with hierarchical clustering (complete-linkage). **b** Heatmap of ***Z*** matrix inferred with MOTL factorization of the target dataset. **c** Reactome enrichment results for direct MOFA and MOTL factors. We performed gene set enrichment analysis on the direct MOFA and the 11 MOTL factors that were differentially active between normal and cancer samples, and that explained at least 1% of mRNA variance. This yielded a set of Reactome processes and pathways significantly associated with either one direct MOFA factor or one or more of 10 different MOTL factors. The Reactome processes and pathways were filtered, clustered, and plotted with orsum. The colors represent the quartiles of enrichment significance for each factor (darker means more significant).

Further statistical tests revealed that only one of the eight factors identified by direct MOFA factorization was differentially active between normal and cancer samples, whereas 19 of the 25 factors obtained by MOTL transfer learning factorization were differentially active (Methods 5.8). Gene set enrichment analysis using Reactome, GO:CC, GO:BP and KEGG (Methods 5.8), focusing on differentially active factors that explained at least 1% of mRNA variance, revealed 715 processes and pathways associated with the direct MOFA factor, and 1061 processes and pathways associated with 11 MOTL factors (Supplementary Table 3). The overlap between the two sets of associated processes and pathways was 318, which is statistically significant (hypergeometric test, one-sided p-value *<* 0.01). We filtered and integrated these enrichment results (Figure 4**c**; Supplementary Figures S3-S5) using orsum (Ozisik et al., 2022) (Methods 5.8). This analysis revealed that MOTL factors capture a broader spectrum of biological pathways than the direct MOFA approach, including for instance immune/inflammatory processes and lipid metabolism pathways.

We also applied both direct MOFA, and MOTL factorizations, to target datasets comprised of normal samples and samples from just a single cancer subtype. In all cases, the direct MOFA factorization yielded only one differentially active factor, whereas MOTL yielded 11 (CL vs normal), 16 (MS vs normal), and 12 (PN vs normal) differentially active factors. Focusing on the subset of differentially active factors that explained at least 1% of mRNA variance, we performed gene set enrichment analyses. We identified processes and pathways that were associated with only a single cancer subtype, such as fatty acid metabolism enrichment for the MS subtype and clathrin-mediated endocytosis for the PN subtype (Supplementary Table 4).

## 3 Discussion

We presented MOTL, which factorizes a multi-omics target dataset by incorporating latent factor values already inferred from the factorization of a multi-omics learning dataset. In the application of MOTL, we are concerned with the situation in which we analyse a target dataset comprised of a limited number of samples. In our evaluations, the target datasets never contained data for more than 15 samples, as our primary concern was with target datasets considered too small for useful factor analysis. It would be insightful to extend the evaluations by using a larger range of sizes for target datasets, in order to identify a crossover point at which transfer learning no longer enhances matrix factorization. In addition, with target datasets containing few samples, the learning dataset is likely to represent a set of biological conditions that neither is entirely specific to, nor fully overlaps with the set of biological conditions represented by the target dataset. It would be relevant to investigate how similar the learning and target datasets need to be in order for informative shared factors to exist. For instance, it would be beneficial to use a measure of similarity, between a given learning and target dataset, which would predict the effectiveness of using a transfer learning approach to apply matrix factorization to the target dataset. It could also be interesting to quantify how heterogeneous (i.e., representing a large diversity of tissues, diseases, experimental conditions …) a learning dataset needs to be, in order to yield factors which can be relevant for a given target dataset.

To evaluate MOTL, we designed two evaluation protocols, based on simulated and real data. Importantly, these protocols can be reused to evaluate other transfer learning strategies for multi-omics data integration. For the evaluation protocol based on real data, we used TCGA, a public repository of multi-omics data with a large number of samples, representing various cancer and tissue types. We selected three different cancer types as references, from which to build target datasets. We did not include samples from these cancer types in the learning dataset. In this setting, it was unknown, prior to evaluation, whether there were latent factors common to the learning and target datasets. Yet, MOTL was effective in uncovering differentially active latent factors, demonstrating that latent factors can be shared across different cancer types. We envisage MOTL as being a helpful tool in the study of rare diseases in general. Therefore a future extension of our work would be to evaluate the application of MOTL to target datasets with non-cancer rare disease samples, using factors inferred from the TCGA learning dataset. We foresee the results of such an evaluation being accompanied by measures of how similar the target datasets are to the TCGA learning dataset, allowing for guidance on when MOTL is a suitable tool for the analysis of other rare disease datasets.

With MOTL we have designed a transfer learning framework that is compatible with a prior learning dataset factorization, as carried out with the MOFA Bayesian approach (Argelaguet et al., 2018). The appeal of a Bayesian framework is the flexibility with regard to the incorporation of prior information, and variational inference serves as a fast alternative to sampling methods. In the future, MOTL could be extended to allow information to be incorporated at other levels of the assumed hierarchy of latent variables. For example, instead of fixing the feature weight values, they could be treated as random variables by MOTL, with priors informed by the factorization of the learning dataset.

In addition to MOFA, there are numerous methods available for multi-omics matrix factorization (Chalise and Fridley, 2017; Lock et al., 2013; Rohart et al., 2017; Mo et al., 2018). A future extension of our work could be a transfer learning framework matched to different multi-omics matrix factorization methods. Similarly, we foresee value in extending an existing transfer learning method that has been designed for single omics data (Sharma et al., 2020; Taroni et al., 2019), so it can be applied in the multi-omics context. The evaluation of such a method, based on the factorization of a learning dataset with a variety of matrix factorization methods, would be informative.

A limitation with MOTL is that we are restricted to a factorization based on features that were retained for factorization of the learning dataset. A consequence is that some features which are highly variable in the target dataset may not contribute to the MOTL factorization. Therefore a future extension could be to add flexibility into the MOTL workflow, so that all features that are highly variable in the target dataset contribute to the factorization, even if they were not retained for the factorization of the learning dataset. Adding flexibility in this way may enhance the weight matrices that are used by MOTL. In the evaluation based on TCGA data, we observed that factors from the groundtruth weight matrices were more easily uncovered using those transferred from the learning dataset than by using those from direct MOFA factorization. However, the transferred factors were slightly less correlated with the factors from the groundtruth weight matrices than the direct MOFA factors were. Despite this, MOTL factorizations were more effective than direct MOFA factorizations in uncovering differentially active groundtruth latent factors. We attribute this superior performance to the fact that the factors in the score matrices inferred by MOTL showed higher correlations with factors from the groundtruth score matrices than the direct MOFA factors. We thus expect that incorporating greater flexibility in the MOTL workflow to retain all of the features that are highly variable in the target dataset would further enhance the weight matrices and produce even more informative factors.

Recently, deep learning, generative, and foundation models have been tested for multiomics data integration (Baião et al., 2025; Ballard et al., 2024; Wen et al., 2023). We however did not identify benchmarks comparing linear methods based on MF with deep learning methods in the context of bulk multi-omics data integration, in particular for small target datasets. It will be interesting to see the developments in this context with more multi-omics data becoming available.

## 4 Conclusions

We presented MOTL, an approach for multi-omics matrix factorization with transfer learning, which infers latent factor values for a multi-omics target dataset comprised of a small number of samples. MOTL factorizes the target dataset by incorporating latent factor values already inferred from the factorization of a learning dataset. We designed two protocols, based on simulated and real multi-omics datasets, for evaluating the performance of multi-omics matrix factorization with transfer learning. We implemented these protocols to evaluate MOTL, and observed that MOTL was more effective in uncovering differentially active groundtruth latent factors than direct matrix factorization without transfer learning. Finally, the application of MOTL to a glioblastoma dataset, comprised of a small number of samples, revealed an enhanced delineation of cancer status and sub-type thanks to transfer learning. We thus demonstrated, in the case of a multi-omics dataset comprised of a small number of samples, that MOTL can enhance the discovery of biological processes and pathways associated with a biological condition of interest. MOTL is accessible as an open source R implementation, as are the evaluation protocols we used in this study.

## 5 Methods

### 5.1 Mathematical Notation

- We denote matrices and datasets with bold capital letters: ***Y***
- If ***Y*** denotes a matrix, we introduce it as ***Y*** = [*y*_*nd*_] ∈ ℝ^*N ×D*^ for which:
- there are *N* rows and *D* columns
- *y*_*nd*_ denotes the value in the *n*th row and the *d*th column
- ***y***_*n*:_ denotes the *n*th row vector, and ***y***_:*d*_ denotes the *d*th column vector
- If ***Y*** denotes a dataset comprised of multiple matrices, we specify this, and denote each of the matrices as 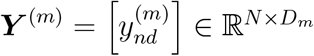
- We denote parameters for statistical distributions as non-bold, lower case letters. If the parameter is for a random variable stored in a matrix, we add indices. For example, 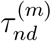 is a parameter for a random variable in the *n*th row and *d*th column of the *m*th matrix of some dataset. If 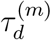 is a parameter for the same matrix, then it is used for all values in the *d*th column.

### 5.2 The MOFA model

Consider a multi-omics dataset ***Y*** consisting of omics matrices ***Y*** ^(*m*)^, *m* = 1, …, *M*. Each 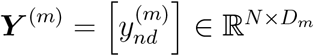 contains data for *N* samples (rows) and *D*_*m*_ features (columns), where 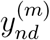 is the value for the *n*th sample and the *d*th feature from the *m*th matrix. MOFA (Argelaguet et al., 2020), assumes the existence of latent factors, and jointly factorizes each ***Y*** ^(*m*)^ into a shared matrix of sample scores ***Z*** = [*z*_*nk*_] ∈ ℝ^*N ×K*^, and an omics specific matrix of feature weights 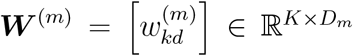. The *k*th column of ***Z*** contains scores for the *k*th factor, and the *k*th row of ***W*** ^(*m*)^ contains corresponding weights for that factor.

MOFA assumes that each observed 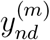 is a random variable, characterised by a probability distribution conditional on a set of latent random variables ***β***. It is assumed that the joint likelihood *p*(***Y*** |***β***) is equal to 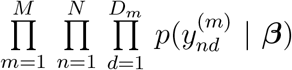, and the choice of probability distribution depends on the type of observed data. A Gaussian likelihood is assumed for observed continuous data:

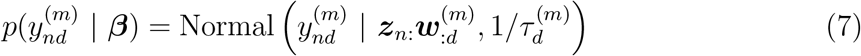

where ***z***_*n*:_ is the vector of scores for the *n*th sample, 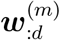 is the vector of weights the *d*th feature from the *m*th matrix, and 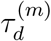 is the precision for that feature. A Bernoulli likelihood is assumed for observed binary data and the logistic link function *π*(*x*) = (1 + *e*^−*x*^)^−1^ is used:

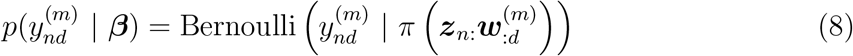

A Poisson likelihood is assumed for observed counts data and the link function *λ*(*x*) = log(1 + *e*^*x*^) is used:

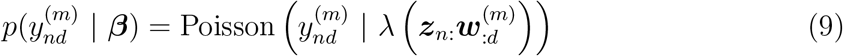

The assumed joint prior distribution, *p*(***β***), is comprised of independent priors: 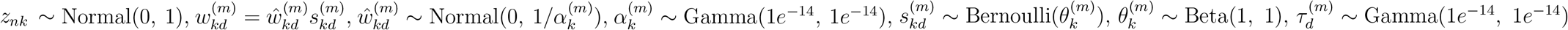.

MOFA uses mean-field variational inference (Grimmer, 2011; Fox and Roberts, 2012; Blei et al., 2017) to approximate the joint posterior distribution, *p*(***β*** | ***Y***), with a joint variational distribution factorized over *J* disjoint groups of variables:

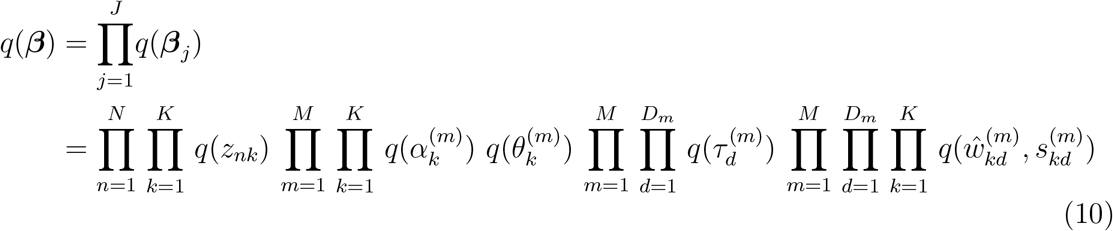

MOFA infers *q*(***β***) iteratively until convergence. At each iteration, each *q*(***β***_*j*_) is updated as

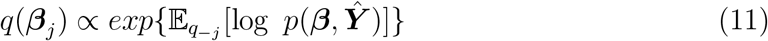

where 𝔼_*q*−*j*_ denotes an expectation with respect to the joint variational distribution, after removing *q*(***β***_*j*_). The dataset ***Ŷ*** is derived by transformation of ***Y***. Observed data with a Gaussian assumed likelihood are transformed with feature-wise centering, which avoids the need to estimate intercepts. Observed data with a non-Gaussian assumed likelihood are transformed to derive Gaussian pseudo-data. The derivation of Gaussian pseudo-data occurs at each iteration, and is based on a new parameter, 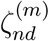, which is derived for each sample *n* and feature *d* from matrix *m*. For observed data with a Bernoulli assumed likelihood, a precision parameter, 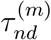, is introduced for each sample and feature as part of the transformation:

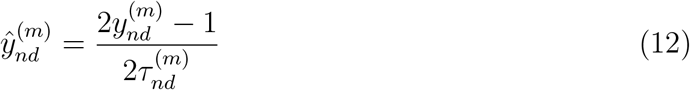

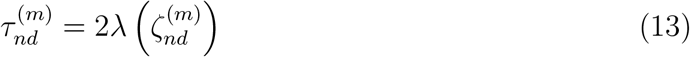

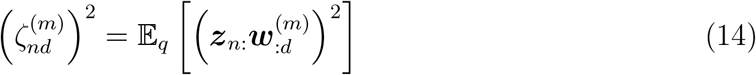

For observed data with a Poisson assumed likelihood, a precision parameter, 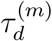, is introduced for each feature as part of the transformation:

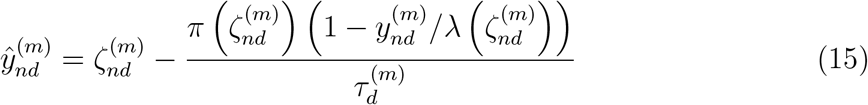

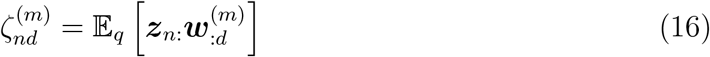

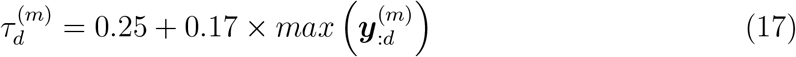

where 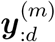 is the vector of observed values for the *d*th feature from the *m*th matrix. For both Bernoulli and Poisson observed data, the 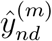 values are centered at each iteration, and 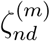 values are derived using the factorization fit from the preceding iteration. MOFA monitors convergence with the evidence lower bound (ELBO), which is used to evaluate how well the variational distribution approximates the posterior distribution. The ELBO is calculated as:

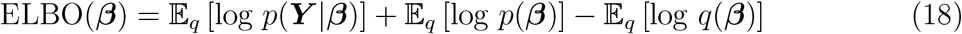

For ***Y*** ^(*m*)^ with non-Gaussian assumed likelihood, MOFA uses a lower bound for each 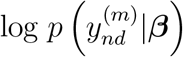. Maximizing this lower bound, coupled with the use of 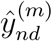 values, allows updates of *q*(***β***) based on the assumption of Gaussian observed data (Jaakkola and Jordan, 2000; Seeger and Bouchard, 2012). MOFA assesses convergence at regular intervals, based on the percentage change in ELBO after. MOFA allows factors to be dropped during training, based on the fraction of variance explained for each matrix. After each iteration, MOFA identifies factors that do not explain a fraction of variance, for any omics matrix, over a threshold. MOFA then drops one of the identified factors.

### 5.3 Multi-omics data simulated with groundtruth factors

We simulated multi-omics datasets, from groundtruth factors, with various configurations. For each simulation configuration we generated 30 instances of a multi-omics dataset, ***Y***, consisting of matrices, ***Y*** ^(*m*)^, *m* = 1, 2, 3. We split each ***Y*** into a target dataset, ***T***, and a learning dataset, ***L***. Each 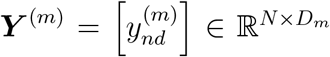 contained data for *N* = *N*_*t*_ + *N*_*l*_ samples (rows) and *D*_*m*_ = 2000 features (columns), where 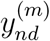 is the value for the *n*th sample and the *d*th feature from the *m*th matrix. *N*_*t*_ is the number of samples subsequently belonging to ***T***, and *N*_*l*_ is the number of samples belonging to ***L***. We generated each ***Y*** ^(*m*)^ from a different statistical distribution, conditional on a random matrix of sample scores, ***Z*** = [*z*_*nk*_] ∈ ℝ^*N ×K*^, and a random matrix of feature weights, 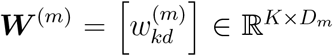. The *k*th column vector of ***Z*** contained sample scores for the *k*th groundtruth factor. The *k*th row vector of ***W*** ^(*m*)^ contained feature weights for that same factor. We varied the number of groundtruth factors across configurations, using *K* ∈ {20, 30}.

We generated ***Z*** based on each sample being a member of a group. In each instance we created two groups of five samples belonging to ***T***, meaning *N*_*t*_ was always equal to 10 samples. We allowed *N*_*l*_ to vary across instances, with samples belonging to ***L*** being in differently sized groups of randomly selected sizes. We used either 20 learning groups of size ∈ {10, 20, 30}, or 40 groups of size ∈ {10, 25, 40}. For the *n*th sample, and *k*th groundtruth factor, we generated the score as *z*_*nk*_ *∼* Normal(*µ*_*g*(*n*)*k*_, *σ*_*z*_), where *µ*_*g*(*n*)*k*_ is the mean parameter for groundtruth factor *k*, for the group that sample *n* belonged to, *g*(*n*). In each instance we selected *µ*_*g*(*n*)*k*_ randomly for each group and groundtruth factor, with probabilities *Pr*(3) = 1*/*8, *Pr*(5) = 3*/*4, *Pr*(7) = 1*/*8. The *k*th groundtruth factor was differentially active for ***T*** if *µ*_*g*(*n*)*k*_ differed between the two target dataset groups. For all instances of a given simulation configuration, the same standard deviation parameter, *σ*_*z*_, was shared by all groups and groundtruth factors. We varied the latent noise-to-signal ratio across our simulation configurations by using *σ*_*z*_ ∈ {0.5, 1.0}

For the *k*th groundtruth factor, and the *d*th feature from the *m*th matrix, we generated the weight as 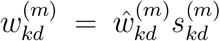. As such, each 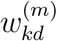 was the product of a continuous random variable, 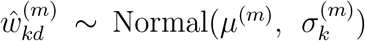, and a binary random variable, 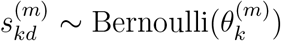. We specified *µ*^(*m*)^, the mean parameter for the *m*th matrix, with *µ*^(1)^ = 5; *µ*^(2)^ = 0; *µ*^(3)^ = 0. We generated, 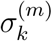, the standard deviation parameter for the *k*th groundtruth factor and the *m*th matrix, with 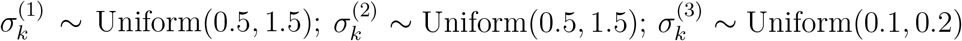. We generated the sparsity for the *k*th groundtruth factor and the *m*th matrix, 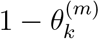, with 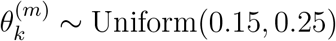.

We generated the values in each ***Y*** ^(*m*)^ as:

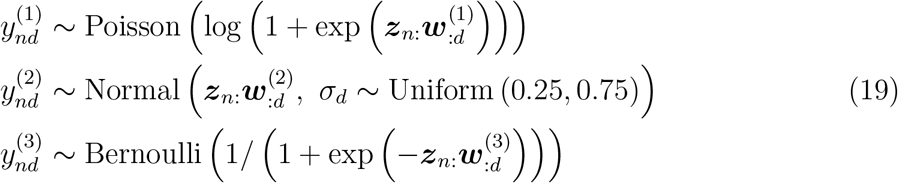

where ***z***_*n*:_ is the vector of scores for the *n*th sample, 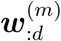 is the vector of weights the *d*th feature from the *m*th matrix, and *σ*_*d*_ is the standard deviation for the *d*th feature.

We split each ***Y*** ^(*m*)^ into 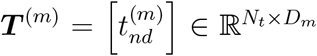, which contained values for the target group samples, and 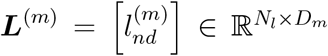, which contained values for the learning group samples.

Before direct factorization with MOFA we pre-processed simulated ***T*** and ***L*** datasets by removing features with 0 variance across samples. Before factorization with transfer learning with MOTL, we pre-processed simulated ***T*** datasets by removing features that had 0 variance across samples or that had been removed from the corresponding ***L*** datasets.

### 5.4 TCGA multi-omics data acquisition and pre-processing

We used the R packages *TCGAbiolinks* (v.2.25.3) and *SummarizedExperiment* (v.1.28.0) to download and save TCGA mRNA expression, miRNA expression, DNA methylation, and simple nucleotide variation (SNV) data (Silva et al., 2016; Hutter and Zenklusen, 2018; Mounir et al., 2019). The mRNA and miRNA expression data consisted of raw counts. The DNA methylation data consisted of CpG site *β*-values, which had been derived from HM450 array intensities with R package *SeSAMe* (v.1.16.0) (Zhou et al., 2018). The SNV data consisted of masked somatic mutation files.

We created four reference datasets, using data from three cancer types; acute myeloid leukemia (LAML), pancreatic adenocarcinoma (PAAD) and skin cutaneous melanoma (SKCM). Each reference dataset, ***R***, contained multi-omics data for all samples from either two, or all three of the cancer types. We did not include SNV data in ***R*** datasets containing LAML samples, due to the sparsity of SNV data for LAML. We only used samples that had data for all omics of interest, and only included one sample per study participant. We thus had multi-omics data for 134 LAML samples, 157 PAAD samples and 435 SKCM samples. We then randomly split each ***R*** into non-overlapping target datasets. Each resulting target dataset, ***T***, contained multi-omics data for five samples per cancer type.

For the evaluation protocol based on TCGA multi-omics data, we merged data from the remaining 29 cancer types into a learning dataset, ***L***. For this ***L*** we only used samples that had data for all four omics, and only included one sample per study participant. This ***L*** contained multi-omics data (mRNA, miRNA, DNA methylation and SNV) for 7,217 samples.

For the application of MOTL to the pd-GBSC target datasets, we created a new learning dataset by merging data from all 32 cancer types. This new learning dataset contained multi-omics data (mRNA, miRNA, DNA methylation and SNV) for 7,866 samples.

Before direct factorization with MOFA, we pre-processed ***R, T*** and ***L*** datasets in the same way. For mRNA data we removed genes that map to the Y chromosome. For both mRNA and miRNA we removed genes if they had a count of zero in *≥*90% of samples, or had zero variance across samples. We normalized mRNA and miRNA counts with the *DESeq2* (v.1.38.0) R package (Love et al., 2014), and log_2_(*x* + 1) transformed the normalized counts. For DNA methylation data, we removed CpG sites that map to the X or Y chromosome, were masked during SeSAMe quality control, had missing values in *≥* 20% of samples, or had zero variance across samples. We converted DNA methylation *β*-values to M-values (Du et al., 2010). We included SNV records whose variant classification was either *Frame Shift Del, Frame Shift Ins, In Frame Del, In Frame Ins, Missense Mutation, Nonsense Mutation, Nonstop Mutation, Splice Site* or *Translation Start Site*. We then created binary SNV matrices aggregated by gene and sample. We removed genes from SNV matrices if the mutation rate across samples was *≤*1%. We filtered all omics to include only the 5,000 most variable features. We did not perform any batch effect correction on ***L*** datasets in order to preserve biological signal (Lee et al., 2020). We checked each ***R*** for batch effects with visualizations of UMAP co-ordinates (McInnes et al., 2020).

We used the R package *uwot* (v.0.1.14) to derive UMAP coordinates from MOFA factorizations, and we did not observe the need to correct ***R*** datasets for batch effects.

Before factorization with transfer learning with MOTL, we pre-processed ***T*** datasets by removing all omics features that had zero variance across samples, or that had been removed from ***L*** during pre-processing. We used *DESeq2* to normalize mRNA and miRNA counts with the geometric means from ***L***, and then log_2_(*x*+1) transformed the normalized counts. We converted DNA methlyation *β*-values to M-values, and converted SNV data to binary matrices after filtering on variant classification, as described previously.

### 5.5 Glioblastoma target dataset acquisition and pre-processing

We created four pd-GBSC target datasets, based on multi-omics profiling conducted by Santamarina-Ojeda et al. for four normal brain samples and nine patient-derived glioblastoma stem cell (pd-GBSC) cultures. The nine cancer samples had been previously classified into three subtypes thanks to transcriptome-based signatures: classical (CL), proneural (PN), and mesenchymal (MS). Each pd-GBSC target dataset contained mRNA expression and DNA methylation data for the four normal brain cortex samples, as well as either all nine cancer samples or just the samples from a subtype.

For factorization with transfer learning with MOTL, the pd-GBSC target datasets initially consisted of mRNA expression raw counts and DNA methylation *β*-values. Before factorization with MOTL, we pre-processed a pd-GBSC target dataset by removing all omics features that had zero variance across samples, or that had been removed from ***L*** during pre-processing. We used *DESeq2* to normalize mRNA counts with the geometric means from ***L***, and then log_2_(*x* + 1) transformed the normalized counts. We converted DNA methlyation *β*-values to M-values.

For direct MOFA factorization, without transfer learning, the pd-GBSC target datasets initially consisted of the same mRNA expression data, but already normalized and transformed by Santamarina-Ojeda et al., and DNA methylation *β*-values. Before direct MOFA factorization, we pre-processed mRNA data by removing genes that map to the Y chromosome, if they had a count of zero in *≥* 90% of samples, or had zero variance across samples. We pre-processed DNA methylation data by removing CpG sites that had missing values in *≥* 20% of samples, or had zero variance across samples. We converted DNA methylation *β*-values to M-values. We filtered both omics to include only the 5,000 most variable features.

### 5.6 Application of MOFA to simulated, TCGA and glioblastoma multi-omics datasets

We factorized simulated target, ***T***, and learning, ***L***, datasets with the MOFA Python implementation *mofapy2* (v.0.6.4). The number of factors we used for each MOFA factorization was equal to the lesser of the number of samples and the number of groundtruth factors that were differentially active when simulating the dataset. The *k*th groundtruth factor was differentially active for a dataset if the mean parameter, *µ*_*g*(*n*)*k*_, for the sample scores for that factor, was not the same for all groups of samples in the dataset. We specified observed data likelihoods corresponding to those used for simulating ***Y*** ^(*m*)^ matrices. We set the maximum number of iterations to 10,000 to ensure convergence.

For the remaining settings we used the *mofapy2* defaults, meaning that all datasets were feature-wise centered during factorization fitting.

We factorized pre-processed TCGA reference, ***R***, target, ***T***, and learning, ***L***, datasets with the MOFA Python implementation *mofapy2* (v.0.7.0). We specified Gaussian as the observed data likelihood for mRNA, miRNA and DNA methylation data, and specified Bernoulli as the likelihood for SNV data. For the ***L*** datasets, we started the factorization with 100 factors and allowed factors to be dropped based on the fraction of variance explained, for which we set the threshold to 0.001. We set the threshold so low in order to retain factors that explained little of the variance in ***L***, yet could be potentially relevant for transfer learning. For ***R*** datasets we also started with 100 factors. For ***T*** datasets, we started with the maximum number of factors allowed by MOFA, which was either 10 factors (two cancer types) or 15 factors (three cancer types). For ***R*** and ***T*** datasets we dropped factors based on a threshold of 0.01, in order to only retain relevant factors. For all TCGA datasets, we set the maximum number of iterations to 10,000, to ensure convergence, and the frequency of convergence checking to five, to ensure that the algorithm had stopped dropping factors before converging.

When saving the factorizations of simulated and TCGA ***L*** datasets, we set the expectations argument to *all*. We did this to ensure that the point estimate for each precision parameter was saved in addition to those that are saved by default.

We factorized the pre-processed pd-GBSC target datasets with the MOFA Python implementation *mofapy2* (v.0.7.0). We specified Gaussian as the observed data likelihood for the mRNA and the DNA methylation data. We started with the maximum allowable number of factors and dropped factors based on a threshold of 0.01.

### 5.7 Application of MOTL to simulated, TCGA and glioblastoma multi-omics datasets

We applied MOTL to simulated, TCGA and pd-GBSC multi-omics target datasets. For each target dataset, we used point estimates of feature weight and precision values saved from the MOFA factorization of the corresponding learning, ***L***, dataset. For observed data with a Gaussian or Poisson assumed likelihood, the transferred value of the precision for each feature, 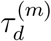, was held fixed throughout iterations of MOTL updates. For observed data with a Bernoulli assumed likelihood, we initialized the value of the precision for each sample and feature, 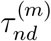, with a feature-wise average, 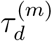, of the 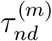 values from the factorization of ***L***. The precisions for Bernoulli observed data were then iteratively updated by MOTL. We estimated intercepts using likelihoods assumed for ***L***, combined with outputs from the MOFA factorization of ***L***. For Gaussian observed data we calculated the intercept for the *d*th feature, from the *m*th matrix, as 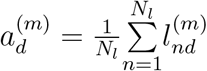, where 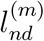 denotes an uncentered learning dataset value after pre-processing. For Poisson and Bernoulli observed data we obtained maximum likelihood estimates of 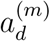 values, for which we used the *mle* function from the R package *stats4* (v.4.2.0). For Poisson observed data we initialized each estimate with 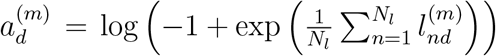, and for Bernoulli observed data we initialized it with 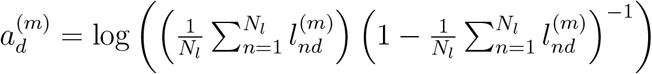.To evaluate robustness, we also applied MOTL to each simulated target dataset after permuting the values in feature vectors of the weight matrices inferred from ***L***. We used a range of proportions for each simulation instance, with the proportions of feature weight vectors permuted in each instance being:

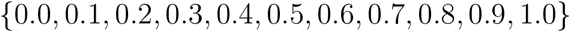

When checking the ELBO for convergence, we used 0.0005% as the threshold, which is the default for MOFA. The algorithm was stopped when the absolute change in ELBO was under this threshold for two consecutive checks, and we set the maximum number of iterations to 10,000 to be consistent with our application of MOFA. For TCGA and pd-GBSC target datasets, and for simulated target datasets factorized after permuting feature weight values, we allowed factors to be dropped based on a threshold of 0.01 for the fraction of variance explained. We checked the ELBO after every five iterations, to ensure that the algorithm had stopped dropping factors before converging.

### 5.8 Evaluation methods

#### Groundtruth factors

For each simulated ***T*** (Methods 5.3), the groundtruth factor values were contained in the corresponding simulated ***Z*** and ***W*** ^(*m*)^ matrices. The sample scores for the *k*th groundtruth factor were contained in ***z***_:*k*_, the *k*th column vector of simulated ***Z***. The feature weights for the *m*th matrix, for that same groundtruth factor, were contained in 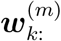, the *k*th row vector of simulated ***W*** ^(*m*)^. For each TCGA ***T*** (Methods 5.4), the groundtruth factors were based on the ***R*** dataset which we had split to create ***T***. We factorized each ***R*** with MOFA, and treated the inferred ***z***_:*k*_ and 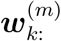 vectors as groundtruth factors for each ***T*** that had been created by splitting ***R***.

#### Differentially active groundtruth factors

For each simulated and TCGA ***T***, groundtruth factor *k* was differentially active if the group means for groundtruth ***z***_:*k*_ differed between the target dataset groups. For each simulated ***T***, this was the group mean, *µ*_*g*(*n*)*k*_, used to simulate groundtruth ***z***_:*k*_. For each TCGA ***T***, the factorization of corresponding ***R*** provided groundtruth ***z***_:*k*_ and 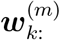 factor vectors. We performed either the Wilcoxian Rank Sum test (two cancer types), or the Kruskal-Wallis test (three cancer types), on each groundtruth ***z***_:*k*_ to determine if there was a statistically significant difference between the cancer types. We classed a groundtruth factor as differentially active if its BH-adjusted p-value was below 0.05.

#### Post-processing

We post-processed inferred and groundtruth ***W*** ^(*m*)^ matrices before evaluation. We scaled each feature vector, 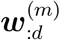, by its Frobenius norm. We then centered each factor vector, 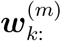, of scaled values separately for each *m*. We then concatenated 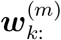 vectors to produce a single vector, ***w***_*k*:_, of centered and scaled feature weights for each factor *k*.

#### Best hits

For each factorization of each simulated and TCGA ***T***, we identified the best hits between the factor vectors inferred with the factorization of ***T***, and the groundtruth factor vectors. For two sets of vectors 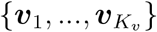 and 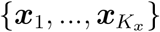 we define the best hit for vector 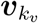 as

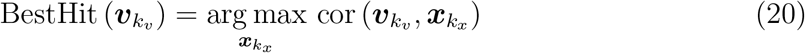

where cor (***v, x***) is the Pearson correlation coefficient between vectors ***v*** and ***x***. We define the best hit for vector 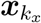 as

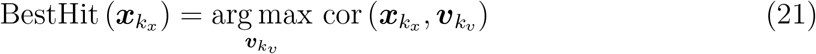

For each simulated ***T***, we identified best hits between inferred and groundtruth ***w***_*k*:_ vectors. For each TCGA ***T***, we identified best hits between inferred and groundtruth ***w***_*k*:_ vectors, as well as between inferred and groundtruth ***z***_:*k*_ vectors. We used shared features when calculating correlations for ***w***_*k*:_ vectors, and we used shared samples for ***z***_:*k*_ vectors. We calculated p-values for the correlations, and only considered correlations with a p-value *<* 0.05 (two-sided alternative hypothesis) when identifying best hits.

#### F-measure values

For each factorization of each TCGA ***T***, we calculated an F-measure value to assess the overall correlation between factor vectors inferred with the factorization of ***T***, and groundtruth factor vectors. We based this on the F-measure presented by Saelens et al., which we adapted in order to assess correlations. For a given set of inferred factor vectors, 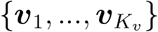, and a set of groundtruth factor vectors, 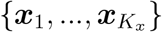, we calculated the F-measure as

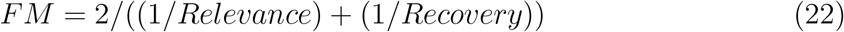

Where

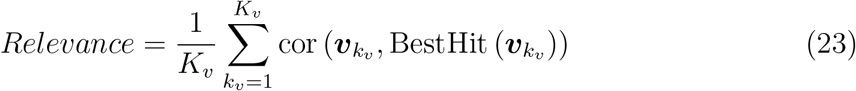

and

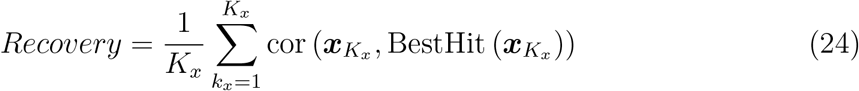

Here cor (***v, x***) is the Pearson correlation coefficient between vectors ***v*** and ***x***. We calculated F-measure values for sets of inferred and groundtruth ***w***_*k*:_ vectors, as well as for ***z***_:*k*_ vectors.

#### F1 scores

We calculated F1 scores to evaluate the factorizations of simulated and TCGA ***T*** datasets:

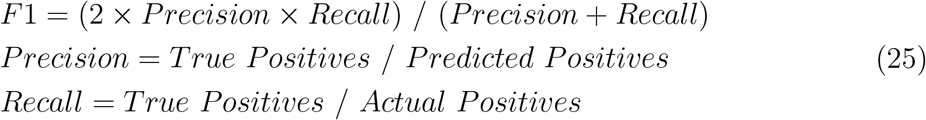

*Actual Positives* were the groundtruth factors of ***T***, that were differentially active.

*Predicted Positives* were the groundtruth factors that were predicted as being differentially active, based on the factorization of ***T***. We firstly performed either the Wilcoxian Rank Sum test (two groups), or the Kruskal-Wallis test (three groups), on each ***z***_:*k*_ vector inferred with the factorization of ***T***, and classed factors with a p-value *<* 0.05 as differentially active. For inferred factors classed as differentially active, we identified the best hits for their corresponding inferred ***w***_*k*:_ vectors. We selected these best hits from the set of groundtruth ***w***_*k*:_ vectors for ***T***. If groundtruth ***w***_*k*:_ was selected as a best hit for a differentially active inferred factor, then groundtruth factor *k* was predicted as being differentially active.

*True Positives* were the differentially active groundtruth factors that were predicted as being differentially active, based on the factorization of ***T***.

#### Statistical testing of differences between factorization methods

We calculated the differences in evaluation measures between factorization methods, and tested the statistical significance of these differences. To do this we fit generalized least squares regressions with the R package *nlme* (v.3.1.157) (Pinheiro and Bates, 2000). We fit a single regression to model the F1 scores for simulated data. For TCGA data we fit a separate regression for each evaluation measure. For each regression we modelled *y*_*i*_ = *β*_0_ + ***d***_*i*_***β***_*d*_ + ***f*** _*i*_***β***_*f*_ + *ϵ*_*i*_. The vector ***d***_*i*_ = (*d*_*i*1_, …, *d*_*iT*_) indicates the simulation configuration, or cancer type, that *y*_*i*_ relates to, and vector ***f*** _*i*_ = (*f*_*i*1_, …, *f*_*iM*_) indicates the factorization method. Vectors ***β***_*d*_ = (*β*_*d*1_, …, *β*_*dT*_)^*⊤*^ and ***β***_*f*_ = (*β*_*f*1_, …, *β*_*fM*_)^*⊤*^ are estimated fixed effects and *ϵ*_*i*_ is the residual. We incorporated correlations between residuals from the same target dataset using the compound symmetry structure method. We calculated contrasts for the factorization method effects in ***β***_*f*_ using the R package *emmeans* (v.1.8.7) (Searle et al., 1980), and used Tukey-adjusted p-values for assessing statistical significance.

#### Differentially active factors from glioblastoma target datasets

We identified differentially active factors from the MOTL factorization of each pd-GBSC target dataset, as well as from the direct MOFA factorization (without transfer learning), of each pd-GBSCs target dataset. We performed the Wilcoxian Rank Sum test on each ***z***_:*k*_ vector inferred with the factorization of a pd-GBSC target dataset. We classed factors with a BH-adjusted p-value *<* 0.05 as differentially active between the normal samples and the cancer samples.

#### Gene set enrichment analysis

We used R package *fgsea* (v.1.24.0) (Sergushichev, 2016) to perform gene set enrichment analysis on differentially active groundtruth factors that were true positives for factorizations of TCGA ***T*** datasets. For each differentially active groundtruth TCGA factor *k*, we analysed vector 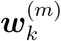 if the fraction of mRNA variance explained by *k* was *>* 0.01, and where *m* corresponded to the mRNA matrix. We tested KEGG, Reactome, GO:BP, and GO:CC gene sets that have a size of between 15 and 500 genes, obtained using the R package *msigdbr* (v.7.5.1). We used an BH-adjusted p-value cutoff of 0.01 for selecting enriched gene sets. We also performed gene set enrichment analysis on differentially active factors from the pd-GBSC target datasets, and used the same criteria as outlined above for differentially active groundtruth TCGA factors. We filtered, clustered, and plotted enrichment analysis results with the Python package *orsum* (v.1.8.0) (Ozisik et al., 2022). We ran orsum with maxRepSize = 2000; maxTermSize = 3000; minTermSize = 15; numberOfTermsToPlot = 30.

### 5.9 Processing time

We used a Dell computer with 20 cores at 3GHz, and 64 GB of RAM, to perform factorizations. To pre-process and factorize the ***L*** used in the TCGA evaluation protocol, it took 26,405 seconds (over seven hours). Hence, we have made the factorization of a large TCGA ***L*** dataset publicly available for transfer learning. It took an average of 37 seconds to pre-process a ***T*** dataset, comprised of four omics, and factorize it directly with MOFA. The average time increased to 134 seconds for MOTL.

## 6 Declarations

### 6.1 Ethics approval and consent to participate

Not applicable.

### 6.2 Consent for publication

Not applicable.

### 6.3 Availability of data and materials

An open source (GPL-3.0 license) R implementation of MOTL is available at https://github.com/david-hirst/MOTL along with the code for the evaluation protocols. The version of the code that can be used to reproduce the analyses in this study is available athttps://doi.org/10.5281/zenodo.15486697. The factorization fit of the full TCGA learning dataset, used for the application of MOTL to the pd-GBSC target dataset, is available at https://zenodo.org/doi/10.5281/zenodo.10847986.

### 6.6 Authors’ contributions

A.B., L.C., P.V., M.V., and D.P.H conceived the project. D.P.H conceived the model and evaluation protocols with input from all authors. D.P.H and M.T implemented the model. D.P.H. generated the figures and results. D.P.H., A.B., and M.V. wrote the manuscript with feedback from all authors. A.B. and M.V. supervised the project. The authors read and approved the final manuscript.

For the purpose of open access, the authors have applied a CC-BY public copyright licence to any Author Accepted Manuscript (AAM) version arising from this submission.

## 6.7 Acknowledgments

We would like to thank Carl Herrmann, Céline Chevalier, Kim-Anh Lê Cao, Lionel Spinelli and Olivia Angelin-Bonnet for helpful feedback and discussions.

