## Supplementary methods and figures for "MOTL: enhancing multi-omics matrix factorization with transfer learning"

### Supplementary methods and figures for the manuscript **MOTL: enhancing multi-omics matrix factorization with transfer learning**

David P. Hirst, Morgane Térézol, Laura Cantini, Paul Villoutreix, Matthieu Vignes, Anaïs Baudot

### 1 Supplementary Methods

#### 1.1 Factorizations with intNMF and moCluster

We factorized simulated and TCGA target, $\boldsymbol{T}$, datasets with intNMF (Chalise and Fridley, 2017) and moCluster (Meng et al., 2016).

##### 1.1.1 Pre-processing

Before factorization of simulated $\boldsymbol{T}$ datasets with intNMF and moCluster, we pre-processed the datasets by removing features with 0 variance across samples. Prior to factorization with intNMF we also forced the data to be non-negative by subtracting the minimum value for each feature vector from all the values in that vector.

Before factorization of TCGA $\boldsymbol{T}$ datasets with intNMF and moCluster, we pre-processed the datasets in the same way as we did before direct factorization with MOFA (Argelaguet et al., 2018): For mRNA data we removed genes that map to the Y chromosome. For both mRNA and miRNA we removed genes if they had a count of zero in $\geq 90\%$ of samples, or had zero variance across samples. We normalized mRNA and miRNA counts with the *DESeq2* (v.1.38.0) R package (Love et al., 2014), and $\log_2(x+1)$ transformed the normalized counts. For DNA methylation data, we removed CpG sites that map to the X or Y chromosome, were masked during SeSAMe quality control, had missing values in $\geq 20\%$ of samples, or had zero variance across samples. We converted DNA methylation $\beta$-values to M-values (Du et al., 2010). We included SNV records whose variant classification was either *Frame_Shift_Del*, *Frame_Shift_Ins*, *In_Frame_Del*, *In_Frame_Ins*, *Missense_Mutation*, *Nonsense_Mutation*, *Nonstop_Mutation*, *Splice_Site* or *Translation_Start_Site*. We then created binary SNV matrices aggregated by gene and sample. We removed genes from SNV matrices if the mutation rate across samples was $\leq 1\%$. We filtered all omics to include only the 5,000 most variable features. Additionally, we had to remove features for which there were missing values, and prior to factorization with intNMF we also forced the data to be non-negative by subtracting the minimum value for each feature vector from all the values in that vector.

##### 1.1.2 Factorization with intNMF

We used the *nmf.mnnals* function from the *InterSIM* (v.2.30.0) R package to apply intNMF to simulated and TCGA $\boldsymbol{T}$ datasets. We weighted each omics matrix as recommended in (Chalise and Fridley, 2017). For simulated $\boldsymbol{T}$ datasets, the number of factors we used for each factorization was equal to the lesser of the number of samples and the number of groundtruth factors that were differentially active when simulating the dataset. For TCGA $\boldsymbol{T}$ datasets, the number of factors we used for each factorization was equal to the number of samples. For all other parameters we used the default settings. It took an average of 762 seconds to factorize a $\boldsymbol{T}$ dataset composed of four omics with intNMF (Dell computer with 20 cores at 3GHz, and 64 GB of RAM).

##### 1.1.3 Factorization with moCluster

We used the *mbpca* function from the *mogsa* R package to apply moCluster to simulated (v.1.42.0) and TCGA (v.1.40.0) $\boldsymbol{T}$ datasets. For simulated $\boldsymbol{T}$ datasets, the number of factors we used for each factorization was equal to the lesser of the number of samples and the number of groundtruth factors that were differentially active when simulating the dataset. For TCGA $\boldsymbol{T}$ datasets, the number of factors we used for each factorization was equal to the number of samples. We specified *method* = *"blockLoading"* and *scale* = *TRUE*. For all other parameters we used the default settings. It took an average of 1 second to factorize a $\boldsymbol{T}$ dataset composed of four omics with moCluster (Dell computer with 20 cores at 3GHz, and 64 GB of RAM).

### 2 Supplementary Figures

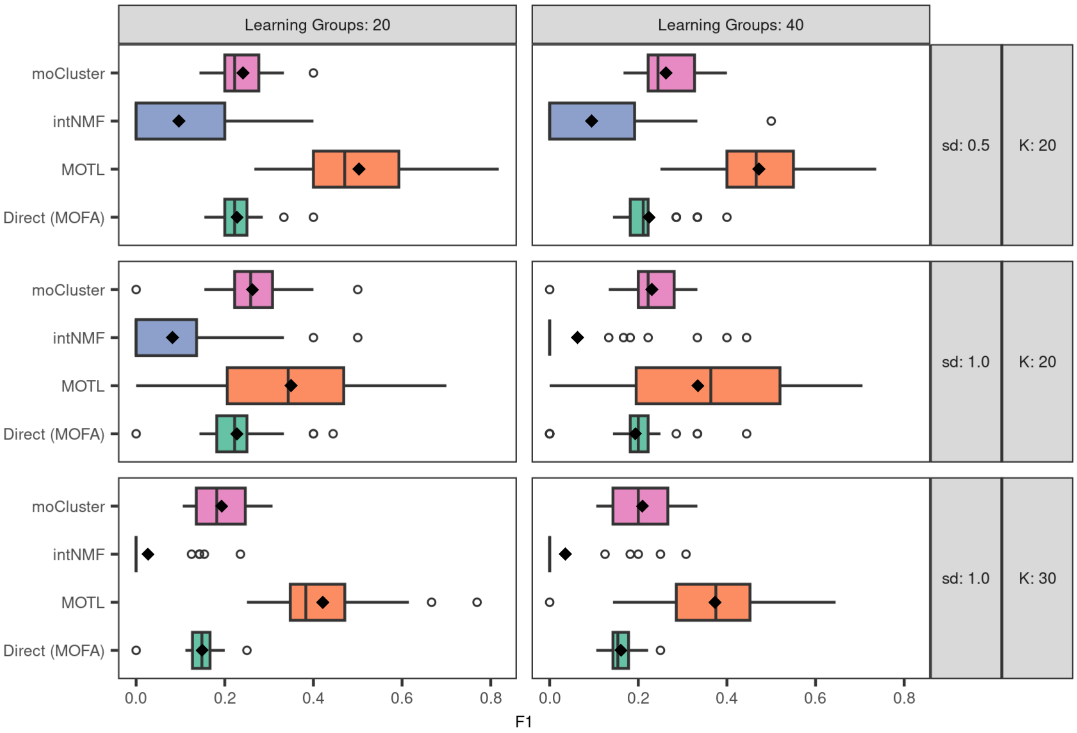


**Figure S1: Evaluation of factorizations of small simulated multi-omics target datasets.** The boxplots represent the F1 scores obtained for factorizations with MOTL transfer learning, MOFA, intNMF, and moCluster, using target datasets generated with different simulation configuration settings. Simulation configurations varied in the number of groups of samples used for the learning dataset (*Learning Groups*), the number of groundtruth factors ($K$), or the standard deviation used to simulate $z_{nk}$ values ($sd$). F1 scores take a value between 0 and 1, and higher values indicate better factorizations. Each boxplot is based on 30 F1 scores. The hinges of the boxes are the $25^{th}$ and $75^{th}$ percentiles, the middle lines are medians, the diamonds are the mean values, and the whiskers are either extreme values or extend 1.5 times the inter-quartile range from the hinge.

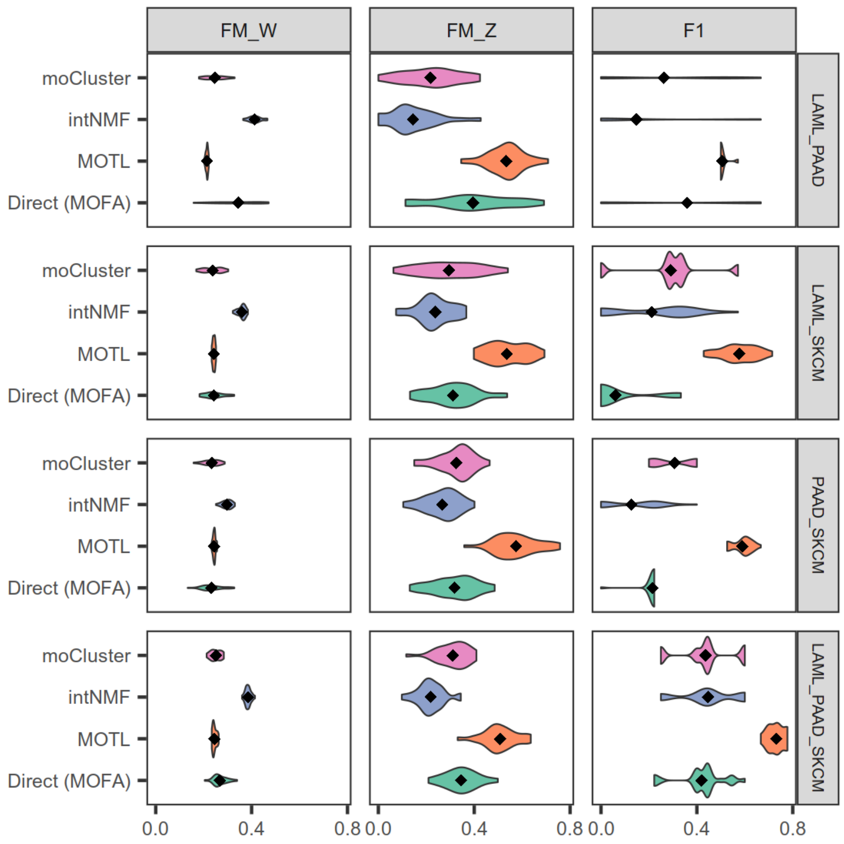


**Figure S2: Comparison of factorization approaches applied to TCGA multi-omics datasets.** Violin plots of F-measure values for weight matrix factors (*FM_W*), F-measure values for score matrix factors (*FM_Z*), and F1 scores (*F1*). For each evaluation score, higher values indicate better factorizations. Scores are plotted by factorization method (MOTL, MOFA, intNMF, moCluster) and by the cancer types characterizing the target dataset samples.

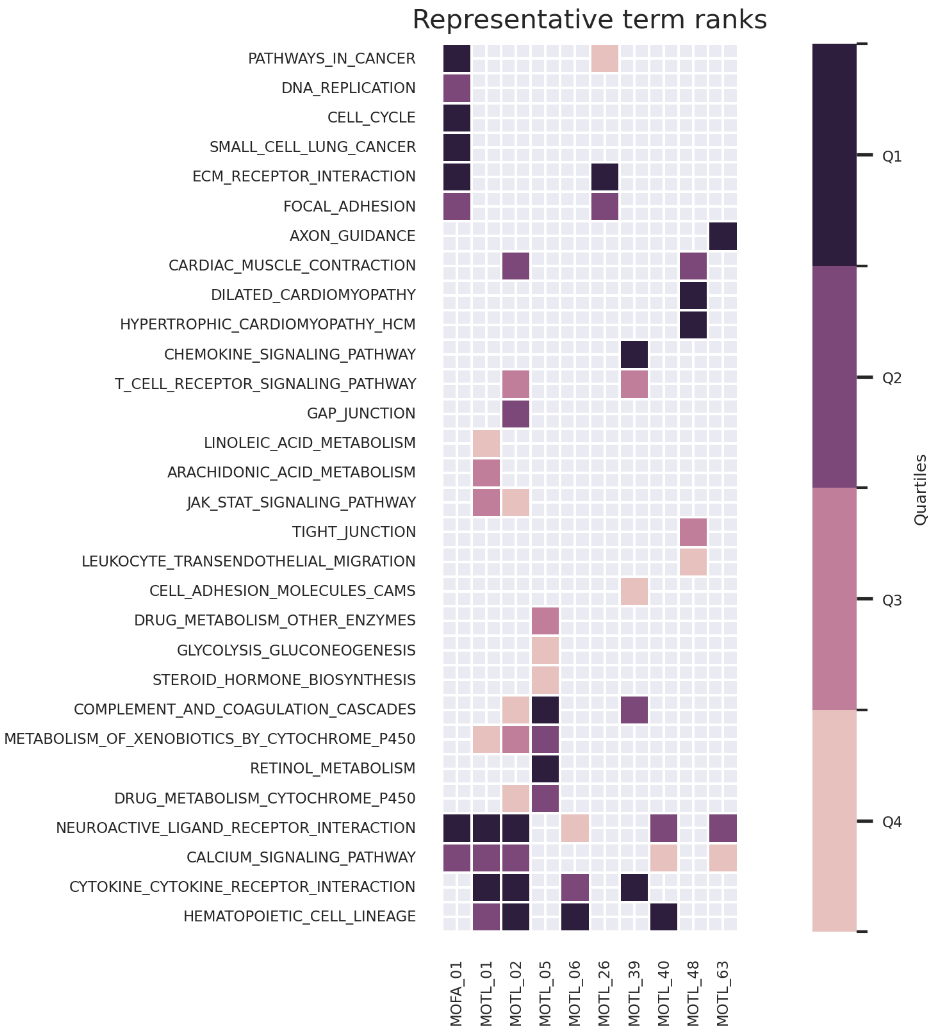


**Figure S3: KEGG enrichment results for direct MOFA and MOTL factors.** We performed gene set enrichment analysis on factors that were differentially active between normal and cancer samples, and that explained at least 1% of mRNA variance. The enriched KEGG processes and pathways were filtered, clustered, and plotted with orsum. The colors represent the quartiles of enrichment significance for each factor (darker means more significant).

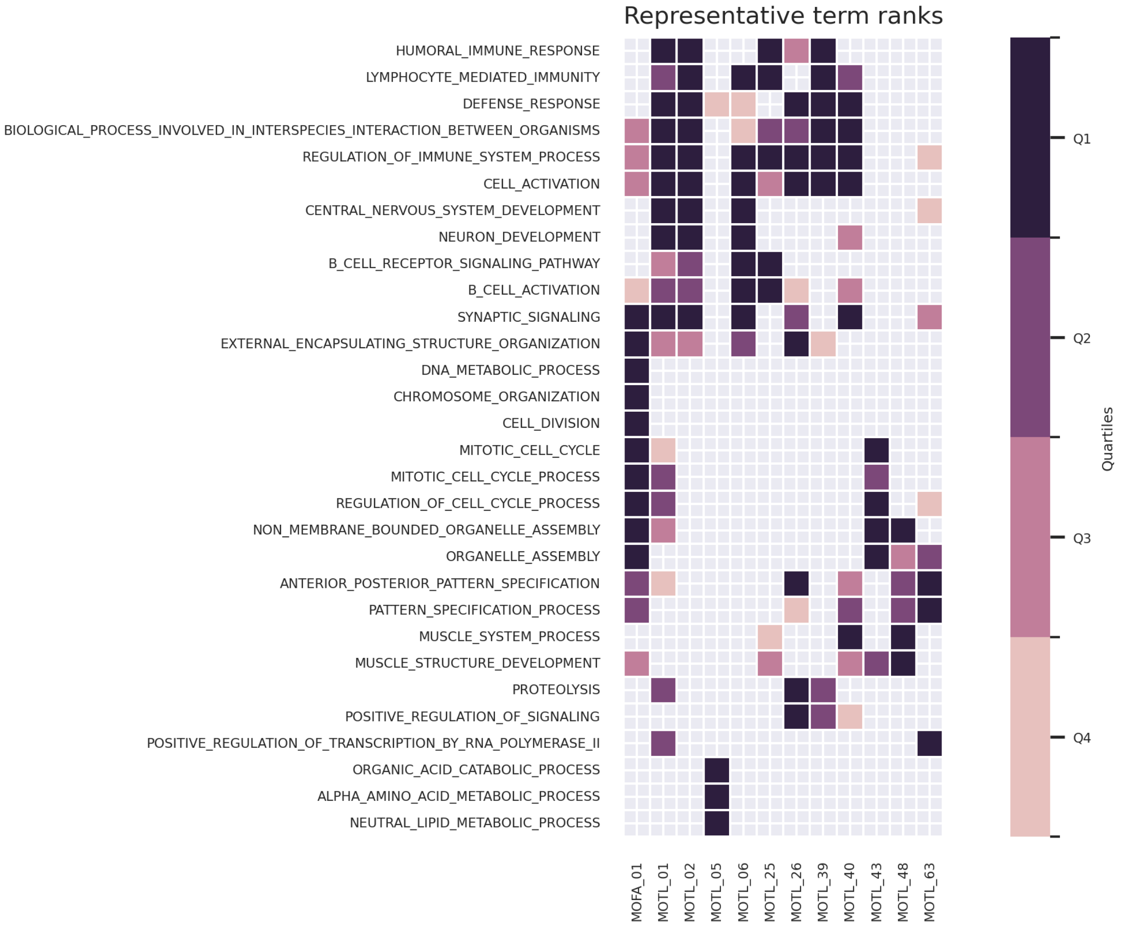


**Figure S4: GO:BP enrichment results for direct MOFA and MOTL factors.** We performed gene set enrichment analysis on factors that were differentially active between normal and cancer samples, and that explained at least 1% of mRNA variance. The enriched GO:BP processes and pathways were filtered, clustered, and plotted with orsum. The colors represent the quartiles of enrichment significance for each factor (darker means more significant).

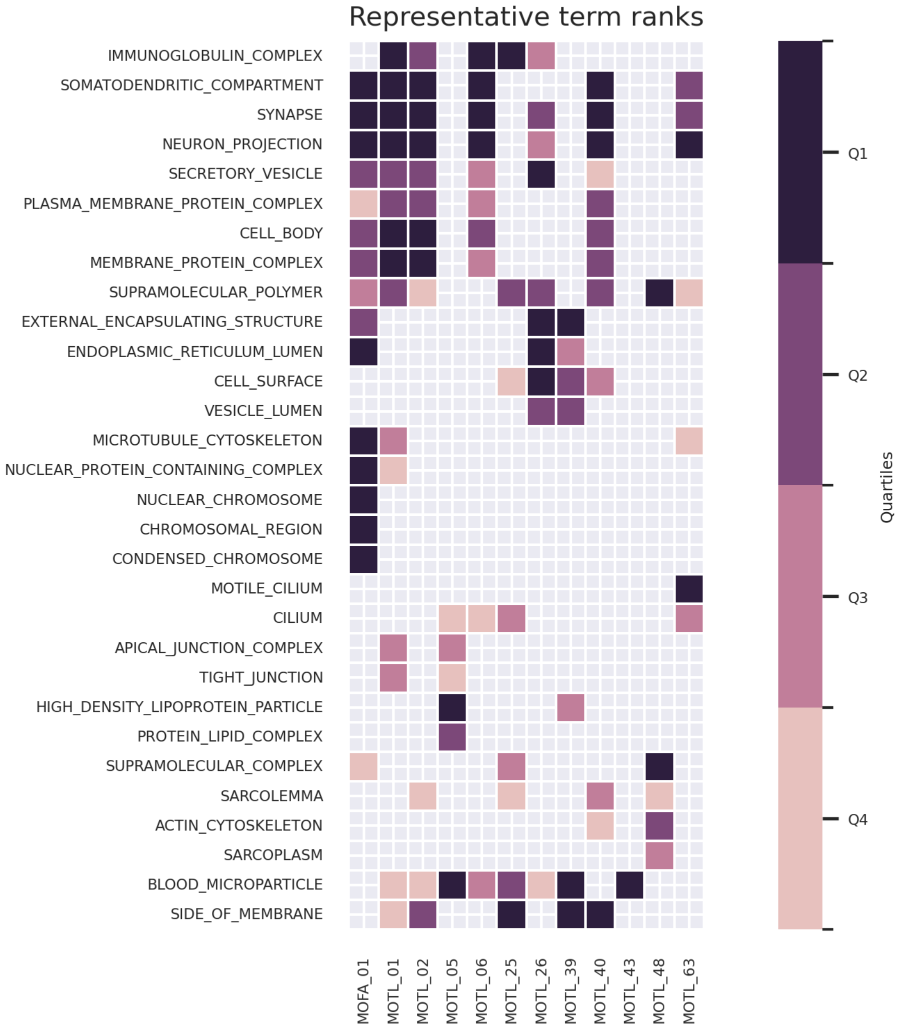


**Figure S5: GO:CC enrichment results for direct MOFA and MOTL factors.** We performed gene set enrichment analysis on factors that were differentially active between normal and cancer samples, and that explained at least 1% of mRNA variance. The enriched GO:CC processes and pathways were filtered, clustered, and plotted with orsum. The colors represent the quartiles of enrichment significance for each factor (darker means more significant).
